# 2C-RT Quantifies Genome-Wide DNA Replication Perturbations and Redefines RIF1–CDC7 Regulation of Replication Timing

**DOI:** 10.64898/2026.10.09.758048

**Authors:** Bruno Urién, Philip Yuk Kwong Yung, Giovanni Giglio, Carmen Navarro, Matilde Fonseca, Sunil Kumar Pradhan, Simon J Elsässer, Bennie Lemmens

## Abstract

Replication timing (RT) coordinates DNA synthesis with genome organization and influences mutation rates, cancer evolution, and developmental plasticity. Current methods infer S-phase timing from DNA content, an assumption that obscures responses to replication perturbations and limits accurate comparisons of RT changes. Here, we develop 2C-RT, a multiplexed sequencing approach that uses DNA content-independent markers for robust RT profiling across S phase. Combining 2C-RT with imaging, we resolve distinct layers of human RT control and identify local and genome-wide responses to drugs and genetic perturbations. We find that RIF1 spatially restricts CDC7 activity and that its loss does not neutralize RT per se but advances replication across most RIF1-sensitive regions. A conserved MCM5 bob1 mutation lowers the origin-activation threshold and reveals a role for CDC7 in establishing local origin dominance. These findings support a revised model in which CDC7 primes early origins, CDK promotes permissive origin firing genome-wide, and RIF1 counteracts CDC7 in distinct chromatin states and peripheral nuclear domains.

## Introduction

In eukaryotic cells, DNA replication starts at multiple sites in the genome following a conserved spatiotemporal program in which distinct genomic regions tend to replicate at defined times during S-phase (Marchal et al., 2019). This replication timing (RT) program is established in early G1 phase and remains stable from one cell cycle to the next, serving as a dynamic yet probabilistic feature of genome duplication with important functional significance (Dimitrova & Gilbert, 1999; Rhind, 2022). The RT program is both influenced by and necessary to maintain chromatin and genome stability, forming an integral feedback loop that sustains cell identity and fitness (Berkemeier et al., 2025; Pope et al., 2014; Klein et al., 2021; Koren, 2014).

Premature replication initiation causes replication-transcription conflicts (Macheret & Halazonetis, 2018; Gómez-González & Aguilera, 2019; Reusswig et al., 2022) and exhaustion of replicating factors (Richards et al., 2022; Zhu et al., 2025). Conversely, delayed replication initiation can compromise complete genome duplication and chromosome segregation in mitosis (Martin et al., 2022; Rainey et al., 2020). Late RT is associated with increased genomic instability, altered gene expression, and epigenetic changes (Donley & Thayer, 2013; Du et al., 2019; Blin et al., 2019). The RT program shapes the distribution of mutations across genomes of multicellular organisms and directly influences the frequency of oncogenic translocations between concurrently replicating regions (Peycheva et al., 2022; Dietzen et al., 2024; Caballero et al., 2023; Weber et al., 2012). Although the relevance of RT is well established, the mechanisms that regulate it are complex and involve multiple layers of temporal and spatial control (Marchal et al., 2019). Hence, accurately detecting RT changes in human cells and determining how different genetic backgrounds and drugs modulate RT remain important challenges in medical biology (Briu et al., 2021).

Multiple methods have been developed to profile RT genome-wide, each relying on the fundamental assumption that S-phase progression correlates with increasing DNA content. Repli-seq sorts asynchronously cycling cells into early- and late S-phase fractions based on DNA content measured by DAPI or PI staining, then label cells with nucleoside analogs such as BrdU or EdU to mark nascent DNA and sequence the replicated regions (Marchal et al., 2019; Rivera-Mulia et al., 2022; Zhao et al., 2020; Wheeler et al., 2025). Alternative single-cell methods use DAPI-based sorting with an RT similarity score (van den Berg et al., 2024) or infer RT from copy number variation in individual S-phase cells, allowing RT profiling without prior cell labeling or sorting (Dashiell J. Massey & Koren, 2022; Massey & Koren, 2022). All these methods implicitly assume that nuclear DNA content reports how long a specific cell has spent in S phase. However, when DNA synthesis is perturbed, DNA content no longer reliably reflects time spent in S-phase or the biochemical state the cell is in. A perturbed cell may have the same DNA content as an unperturbed cell at a much earlier time point, leading to misclassification of S-phase states and masking changes in RT. We anticipate that many RT regulators will affect DNA synthesis and, consequently, the accumulation of DNA content, increasing the risk of bias in classic RT estimates.

We propose that the widespread use of DNA content as a proxy for S-phase time has contributed to several unresolved paradoxes in the field. Central to RT regulation are the loading and phosphorylation dynamics of MCM (minichromosome maintenance) helicases, which form the core of the replication fork. Molecular studies have identified RIF1 (Rap1-interacting factor 1) as a key regulator of RT that acts locally to delay replication initiation by promoting MCM dephosphorylation (Hiraga et al., 2014; Alver et al., 2017). However, RT datasets based on the classic DNA accumulation assumption reveal unexpectedly ambiguous consequences of RIF1 loss. For instance, RIF1 deficiency leads to a flattening of RT profiles rather than causing a general advance in RT, with most genomic regions converging toward mid-S-phase timing (Foti et al., 2016; Klein et al., 2021). A similar inconsistency can be observed for CDC7 (cell division cycle 7-related protein kinase), which is expected to promote early RT through direct phosphorylation of MCM, in opposition to RIF1. Contrary to this prediction, RT-sequencing studies using established CDC7 inhibitors report only modest genome-wide effects, with more than one-third of observed changes corresponding to RT advances when inhibiting CDC7 (Göder et al., 2024). This paradox is further compounded by DAPI-based microscopy studies suggesting a predominant role for CDC7 in late S-phase (Göder et al., 2024; Rainey et al., 2020), whereas genetic and biochemical evidence supports a more critical function during early S-phase, when compensatory CDK (cyclin-dependent kinase) activity remains low (Suski et al., 2022; Spencer et al., 2013; Lemmens et al., 2018; Saleh et al., 2022; Poh et al., 2014). Taken together, these discrepancies, both within and across experimental approaches, underscore a fundamental limitation in current RT measurements and highlight the need for DNA content–independent RT profiling.

Cell-cycle proteins such as cyclins often show phase-specific accumulation patterns and can be used to infer temporal progression from fixed-cell snapshots (Akopyan et al., 2014; Gookin et al., 2017; Stallaert et al., 2022). Notably, Cyclin A2 accumulates progressively during S- and G2-phase and gets degraded by the APC/C (Anaphase-Promoting Complex/Cyclosome) upon mitotic exit (Akopyan et al., 2014; Gookin et al., 2017). Although increased DAPI signal depends on DNA replication, Cyclin A2 levels continue to accumulate when DNA synthesis is abolished (Lemmens et al., 2018; Sosenko Piscitello et al., 2025), making Cyclin A2 a potentially superior marker of S-phase progression under perturbed conditions.

Here, we develop 2C-RT, a nascent-DNA sequencing approach that exploits the cell cycle “to see” RT. 2C-RT uses DNA content-independent timers such as Cyclin A2 to enable multiplexed comparisons of RT across conditions. Combining 2C-RT with time-resolved microscopy, we resolve distinct layers of RT control, including local and genome-wide effects, and redefine the mechanistic relationships among RIF1, CDC7, and MCM5 in establishing the RT program in human cells.

## Results

### Cyclin A2 reveals replication delays masked by DNA content analysis

We compared markers of S-phase progression in asynchronous hTERT-RPE-1 cells (hereafter RPE) using quantitative image-based cytometry (QIBC). Plotting DNA synthesis, based on EdU incorporation, against DNA content produced a characteristic arc-shaped distribution, with distinct G1, S, and G2 populations ordered along a temporal axis. A similar arc-shaped pattern was observed when plotting EdU incorporation against nuclear Cyclin A2 intensity (Figure 1A). The EdU-positive population spanned early S-phase, characterized by low DAPI and low Cyclin A2 levels, to late S-phase, where both signals were high. To assess how delayed DNA replication affects the S-phase population, cells were treated with the CDC7 inhibitor XL413 for 4 hours. This treatment impairs phosphorylation of MCM2 and MCM4, thereby slowing replication initiation across the genome (Alver et al., 2017; Lemmens et al., 2018). XL413 treatment significantly reduced the percentage of EdU-positive cells (Figure S1A) and markedly decreased the EdU intensity in replicating cells (Figure S1B), confirming inhibition of DNA replication and consistent identification of S-phase cells. However, the inferred S-phase timing differed between the two approaches. Cyclin A2-based analysis revealed a clear shift towards high Cyclin A2 intensity within the S-phase population following XL413 treatment, consistent with a pronounced inhibition of early S-phase firing. In contrast, CDC7 inhibition did not clearly shift DAPI intensity distribution compared to untreated EdU-positive cells, suggesting no apparent S-phase delay (Figure 1A and Figure S1C). Similar results were observed across multiple cell lines (RPE, U2OS, BJ). Together, these data indicate that DNA content–based markers such as DAPI poorly capture genome-wide replication delays, whereas Cyclin A2 retains temporal resolution under perturbed conditions (Figure 1B).

**Figure 1.**
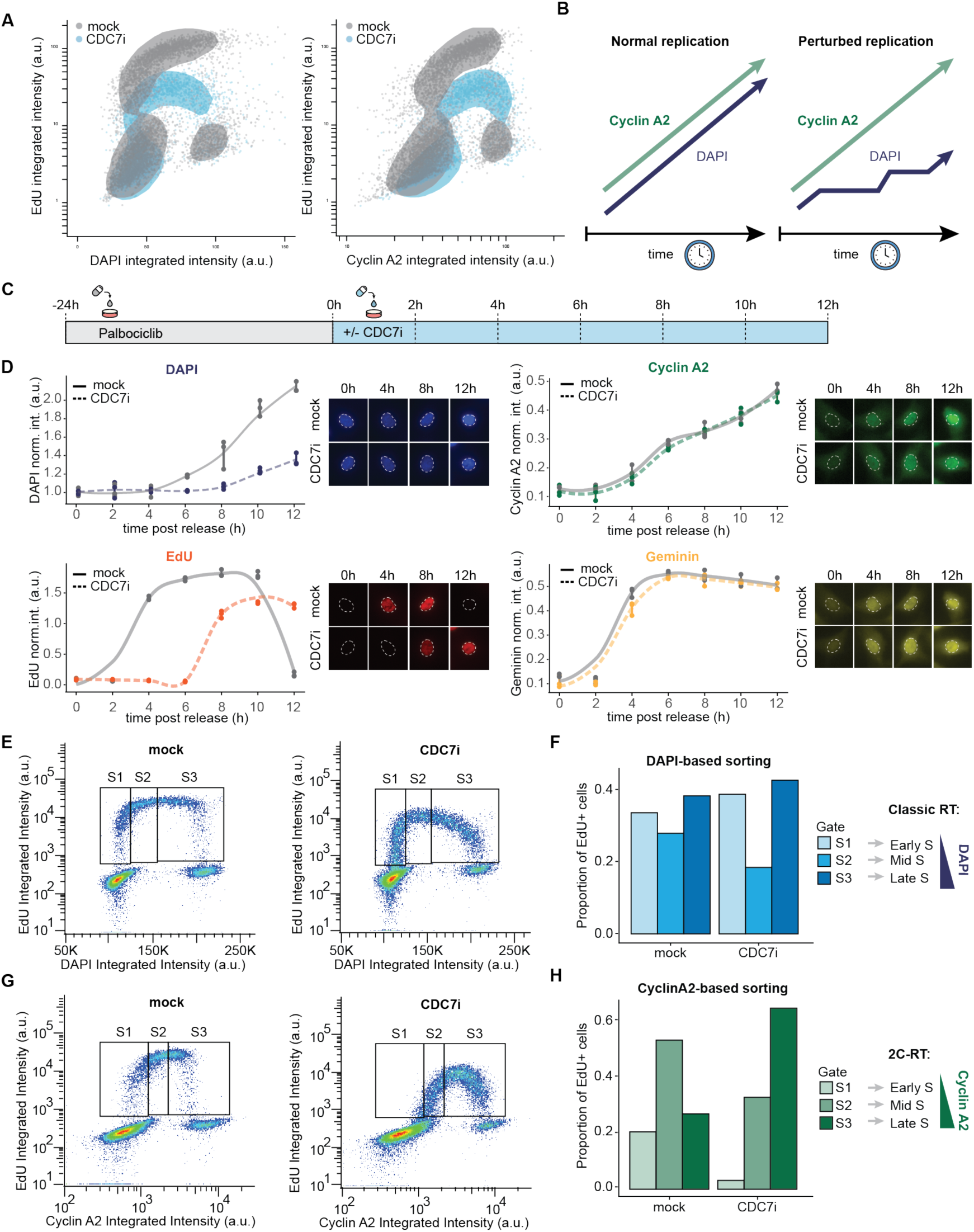
Cyclin A2 is a robust marker for S-phase progression independent of DNA accumulation. (A) Nuclear QIBC signal of RPE cells treated with no drug or XL413 15 μM (Cdc7i) for 4 hours and labelled for nascent DNA with EdU 10 μM in the last 30 minutes prior to fixation. EdU is compared to DNA content (left) and Cyclin A2 accumulation (right). 2D-density area of 80% most dense data points highlighted in the background. (B) Schematic of the expected accumulation of DNA content and Cyclin A2 over time from S-phase onset in cells undergoing normal or perturbed DNA replication. (C) Schematic of time-course experiment setup, consisting of 24 hour G1 synchronization with Palbociclib 200 nM, followed by release in no drug or XL413 15 μM (Cdc7i) for the indicated timepoints and DNA synthesis labelling in the last 30 minutes with EdU 10 μM. (D) Normalized intensity over the time-course of the maximum density signal in RPE cells for the following S-phase markers upon fixation at the given timepoint: DNA content as DAPI (top-left), DNA synthesis as EdU incorporation in the last 30 minutes of the timepoint (bottom-left), Geminin (top-right) and Cyclin A2 (bottom-right). Per timepoint, N = 3. Exemplary images of nuclei in representative timepoints next to each corresponding graph. (E) Flow cytometry data of RPE cells treated with no drug or XL413 15 μM (Cdc7i) for 4 hours and labelled for nascent DNA with EdU 10 μM in the last 30 minutes prior to collection. DNA content reflected as DAPI plotted against EdU incorporated. Non-overlapping gates S1, S2 and S3 cover the range of EdU-positive cells. (F) Proportion of EdU-positive cells in each gate depicted in (E) divided by total number of EdU-positive cells. (G) As in (E), but plotting Cyclin A2 levels against incorporated EdU instead and a different set of non-overlapping gates. (H) Same as (F), but for gates depicted in (G).

To further validate Cyclin A2 as an S-phase timer, we performed a time-course experiment in which RPE cells were synchronized in G1-phase with Palbociclib and released for 2, 4, 6, 8, 10, or 12 hours in the presence or absence of XL413. Cells were fixed after the indicated timepoint and analyzed by QIBC for nuclear DAPI, Cyclin A2, Geminin, and EdU levels (Figure 1D). In control cells, all parameters increased over time, consistent with S-phase progression. Upon XL413 treatment, Cyclin A2 and Geminin levels followed kinetics similar to control cells, whereas EdU incorporation and the increase in DNA content were delayed (Figure 1D). These observations are consistent with previous findings showing that Cyclin A2 accumulation proceeds independently of DNA replication and is maintained upon genetic ablation of essential replication factors (Lemmens et al., 2018; Sosenko Piscitello et al., 2025). Notably, the monotonic increase in Cyclin A2 levels throughout S and G2 phases supports its use as a timer of replication progression, whereas the narrow window during which endogenous Geminin accumulates to maximal nuclear levels may make it more suitable for measurements in early S phase. To use Cyclin A2 for RT studies, defined S-phase fractions must be isolated for downstream analysis, typically by fluorescence-activated cell sorting (FACS). FACS of mock- and XL413-treated RPE cells revealed a shift towards late S-phase fractions when cells were gated according to Cyclin A2 levels, but not when DAPI-based gating was used (Figure 1E-H). Together, these data demonstrate that endogenous Cyclin A2 enables robust isolation of early, mid, and late S-phase populations and reveals global replication perturbations that are masked by traditional DNA content–based classification.

### Development and validation of 2C-RT

Motivated by the discrepancy between DAPI- and Cyclin A2–based cell cycle classification (Figure 1), we developed a genome-wide RT profiling strategy based on Cyclin A2–guided sorting. We termed this approach 2C-RT as it leverages the Cell Cycle to see RT (Figure 2A). The workflow consists of four steps: (1) FACS-based isolation of early, mid, and late S-phase cells using EdU incorporation and Cyclin A2 levels; (2) sample barcoding and pooling via MINUTE-ChIP (Kumar & Elsässer, 2019) for multiplexed library preparation; (3) sequencing of EdU-labeled DNA to generate genome-wide replication profiles; and (4) computation of Replication Timing Index (RTI), as previously described (Van Rechem et al., 2021). RTI summarizes the proportion of normalized reads per sorted gate for a given genomic region, providing a quantitative and easy to read measurement of RT for more than two sorted S-phase fractions (Figure 2A).

**Figure 2.**
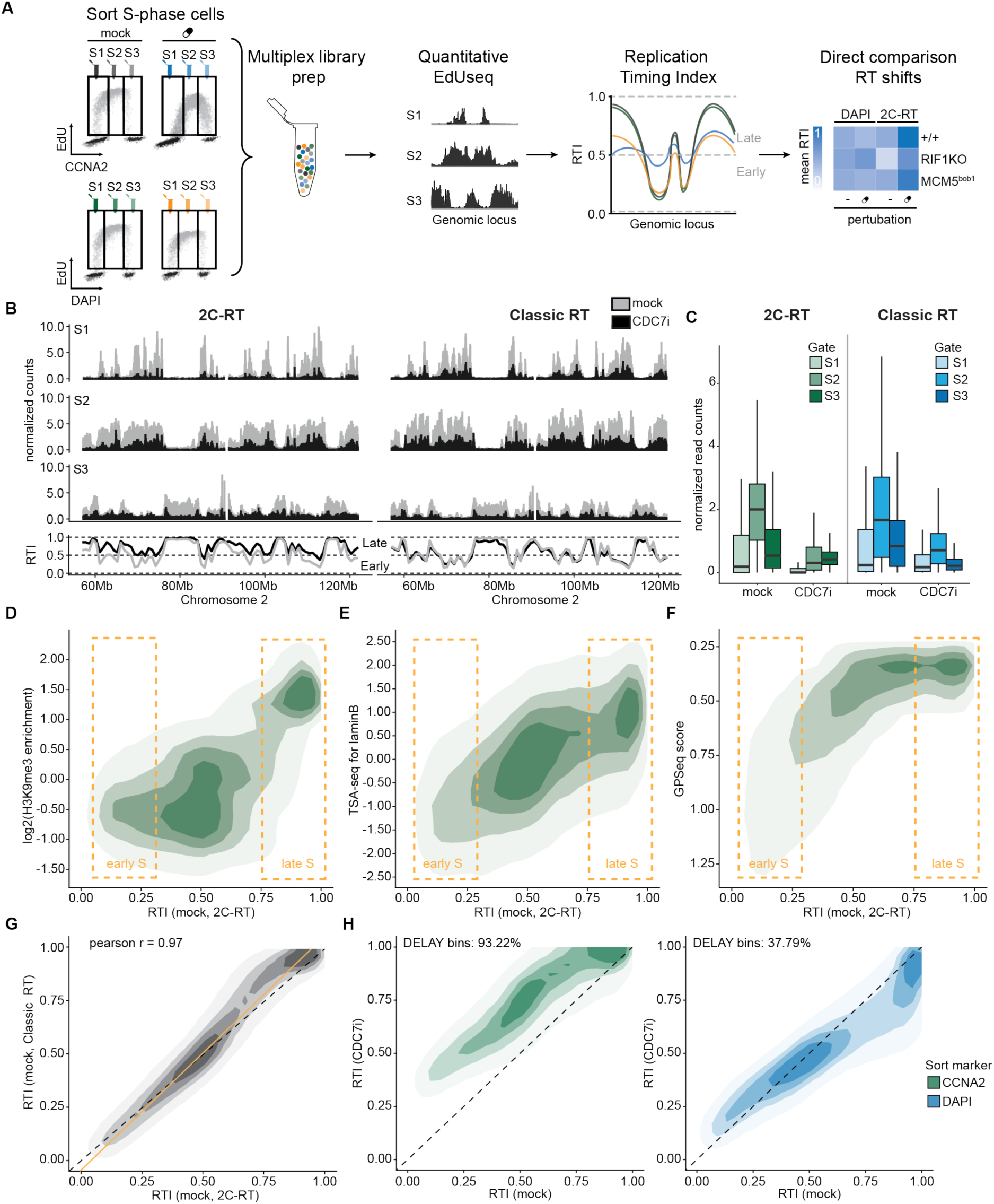
Cell-cycle-derived replication timing properly captures genome-wide delay due to Cdc7 inhibition. (A) Schematic of 2C-RT strategy, including cell sorting with both marker and EdU signal to collect specifically actively replicating cells in different stages of marker signal, followed by multiplex library preparation in which EdU-containing DNA fragments are isolated, sequencing, normalization to obtain RTI per genomic bin and condition, and final downstream analysis like correlation between conditions. (B) RPE cells were treated with no drug or XL413 15 μM (Cdc7i) for 4 hours, labelled with EdU 10 μM in the last 30 minutes of the treatment and processed with the sorting and multiplexed sequency strategy depicted in (A), one replicate per sort. A representative region of human chromosome 2, 57 to 122 Mb in human genome assembly 38, was selected to show the pattern of input-normalized 50-kb-binned reads in each gate (top) and resulting RTI calculated and smoothed by LOESS non-parametric regression (bottom). Left side depicts results of our 2C-RT strategy based on Cyclin A2 staining, whereas right side shows data of traditional Repli-seq strategy based on DAPI staining. (C) Box plots showing normalized read counts per condition and grouped in 50-kb bins (N = 3). In each box, the line within the box indicates the median, and the box edges show the 25th and 75th percentiles. Whiskers extend beyond box limits 1.5 times the interquartile range between 25th and 75th percentiles. (D) 2D-density plot of RTI at 50-kb resolution from our unperturbed 2C-RT data against H3K9me3 enrichment data in RPE cells previously published. Each contour area represents (from lighter to darker) 95%, 75%, 50% and 25% of the most densely regions covered by datapoints in the graph Yellow boxes indicate plot area containing predominantly early or late 50-kb bins based on RTI value (RTI < 0.25 for early S-replicated bins, indicating >50% input-normalized reads in those bins correspond to S1 fraction; RTI > 0.75 for late S replicated bins, >50% input-normalized reads in those bins correspond to S3 fraction). (E) Same as (D), but plotting our unperturbed 2C-RT data against published TSA-seq chromatin proximity data to lamin B1 in RPE cells. (F) Like (D), but plotting our control 2C-RT data against a public GPSeq dataset for RPE cells measuring relative distance to nuclear envelope. (G) Pairwise comparison of each 50-kb bin RTI value in mock 2C-RT data against mock classic RT data. Each contour area represents (from lighter to darker) 95%, 75%, 50% and 25% of the most densely regions covered by datapoints in the graph. Linear model fit in yellow, with pearson correlation indicated at the top left. (H) Pairwise comparison of each 50-kb bin RTI value in mock versus XL413-treated RPE cells. Each strategy is represented separately (2C-RT on the left, traditional Repli-seq on the right). Each contour area represents (from lighter to darker) 95%, 75%, 50% and 25% of the most densely regions covered by datapoints in the graph. Diagonal black line represents the expected location of a perfectly identical RT profile between condition. Percentage of 50-kb bins above the diagonal indicated at the top of each graph as “DELAY”.

We applied both 2C-RT and DNA content-based RT profiling (referred to as “classic” RT profiling here) to RPE p53KO cells treated with either mock or XL413, using matched gating strategies (Figure 1E and 1G). Reads were aligned, input-normalized and aggregated into 50-kb bins for subsequent RTI calculation (Figure 2B). Normalized read distributions per gate were similar in 2C-RT and classic RT strategies, with S2 mid gate accumulating the most reads, followed by S3 and finally S1 (Figure 2C). Consistent with reduced EdU incorporation observed by QIBC and FACS, CDC7 inhibition decreased normalized read counts per bin across all fractions, irrespective of sorting strategy. However, distribution of normalized reads of XL413-treated cells shifted towards gate S3 in 2C-RT, whereas S2 remained the dominant fraction in classic RT (Figure 2B-C).

We compared RTI profiles with independent datasets of chromatin state and nuclear organization. Late-replicating regions (RTI > 0.75) correlated with H3K9me3-enriched heterochromatin (log2 H3K9me3 > 1; Figure 2D). Late-replicating regions also correlated with two spatial features: proximity to nuclear lamina (Lamin B TSA-seq > 0; Figure 2E) and peripheral nuclear positioning (GPSeq < 0.5; Figure 2F). These associations are consistent with established relationships between RT and 3D genome organization (Chen et al., 2018; Girelli et al., 2020; Wang et al., 2021; Yip et al., 2026), supporting the accuracy of 2C-RT.

Pairwise comparison of RTI values obtained by 2C-RT and classic RT profiling showed strong agreement between the two approaches, indicating that both capture steady-state RT similarly (Pearson r = 0.97; Figure 2G). We also applied the 2C-RT strategy to p53KO RPE cells expressing the PIP-FUCCI system (Grant et al., 2018), using exogenous Gem_1-110_-mCherry expression as the S-phase timer (Figure S4E-F). RT profiling based on Gem_1-110_-mCherry levels highly correlated to Cyclin-A2-based 2C-RT, but the former became flatter from mid S-phase progression onwards. The strong ectopic promoter and diminished proteasomal degradation of the Gem_1-110_-mCherry peptide likely contribute to the rapid saturation of marker expression, matching published data of Gem_1-110_-mCherry based reporters (Grant et al., 2018) and the rapid induction of endogenous Geminin upon S-phase entry (Figure 1D). These data further validate the 2C-RT methodology and suggest that Geminin-based timings are confined to early S-phase quantifications.

We next evaluated the similarity between classic RT and 2C-RT under suppressed origin firing. In the 2C-RT datasets, CDC7 inhibition with XL413 delayed replication in the majority of genomic bins (93.22%), with the strongest effects in early-S and mid-S phase regions. In contrast, classic RT indicated delays in only 37.79% of bins and suggested a relative advance of late-replicating regions (Figure 2H). Thus, while both methods agree under unchallenged conditions, they diverge substantially following replication perturbation. Notably, 2C-RT revealed a uniform delay, consistent with the microscopy and flow cytometry data we obtained (Figure 1 and supplemental figures S1 and S3) and with the established role of CDC7 in promoting replication initiation in early S-phase (Poh et al., 2014).

### 2C-RT detects genome-wide and local changes in DNA replication dynamics

We next asked whether 2C-RT could also detect replication advances. In yeast, Cdc7 deficiency can be suppressed by Rif1 loss or Mcm5-bob1 mutation, suggesting that these mutations enhance origin activation when Cdc7 activity is low (Hoang et al., 2007). In mammalian cells, RIF1 also antagonizes CDC7 via its interaction with the PP1 phosphatase, whereas the relevance of the bob1 mutant remains unknown. To model these conditions, we used an established RIF1 knockout line (Chiang et al., 2016; Hawgood et al., 2025) and generated a homozygous MCM5^bob1^ mutation in p53KO RPE cells, corresponding to the yeast Mcm5-bob1 mutant (Figure 3A and Figure S2A). Through CRISPR-mediated gene-editing, we changed MCM5 at the highly conserved proline in position 92 to leucine (P92L), which is structurally equivalent to Mcm5-bob1 P83L mutation in *S. cerevisiae* (Figure S2B).

**Figure 3.**
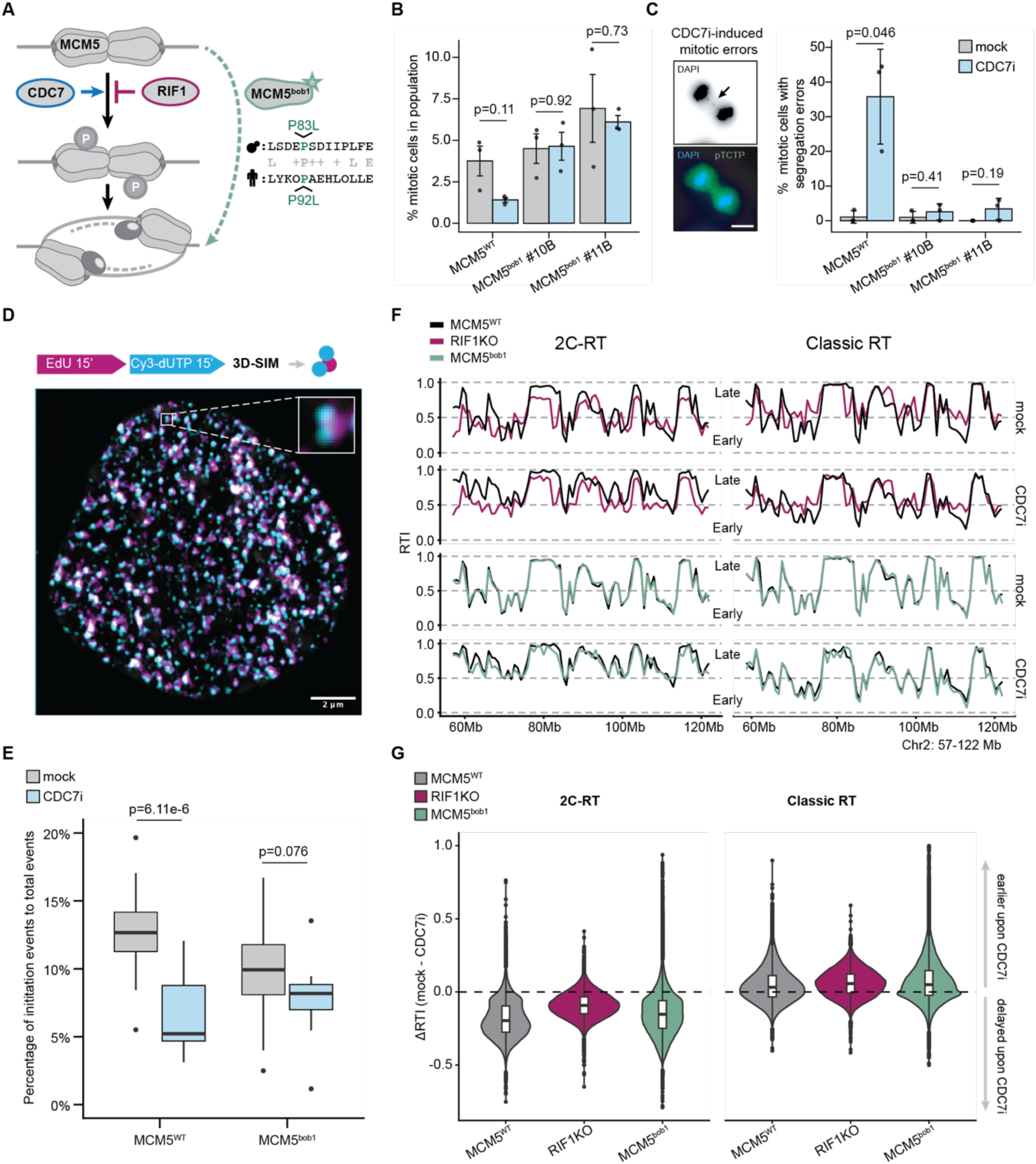
2C-RT retains genome-wide fluctuations of replication timing. (A) Schematic of mechanisms to bypass CDC7 phosphorylation in cells. For MCM5bob1 mutant, a simplification of the mutated proline in *Saccharomyces cerevisiae* and *Homo sapiens* is included at the bottom right. (B) Bar plot of percentage of mitotic cells in RPE p53KO cells containing endogenous MCM5 wild-type or bob1 mutated, N = 3. Each cell line was left treated with no inhibitor or XL413 15 μM (CDC7i) for 8 hours prior to fixation and staining against pTCTP to help identifying mitotic cells under the microscope. Two independent homozygous clones are included. Error bars represent standard deviation of the samples in each condition. Statistical p-values of t-tests indicated between corresponding conditions compared. (C) Representative images (left) and bar plot (right) of mitotic errors within the mitotic population of same experiment setup as (B), N =3. Error bars represent standard deviation of the samples in each condition. Statistical p-values of t-tests indicated between corresponding conditions compared. (D) Representative image of a nucleus dual pulsed for DNA synthesis as in 3D-SPARK protocol. Magenta represents first EdU 10 μM 15 min pulse, cyan represents second Cy3-dUTP transfection pulse. Zoom-in to an example of how an initiation event looks like, with one EdU region progressing into two independent Cy3-dUTP regions. (E) Box plot of percentage of initiation events compared to all dual-pulsed events quantified per nucleus. RPE p53KO cells containing MCM5 wild-type or bob1 mutation were treated with no inhibitor or XL413 15 μM (CDC7i) for 4 hours prior to 3D-SPARK-based labelling, fixation and imaging via 3D-SIM. N =16 nuclei for all conditions except for MCM5bob1 #11B, which was N =14. In each box, the line within the box indicates the median, and the box edges show the 25th and 75th percentiles. Whiskers extend beyond box limits 1.5 times the interquartile range between 25th and 75th percentiles. Dots indicate outliers past the whisker range. Statistical p-values of t-tests indicated between corresponding conditions compared. (F) RPE cells of the indicated genotypes were treated with no drug or XL413 15 μM (Cdc7i) for 4 hours, labelled with EdU 10 μM in the last 30 minutes of the treatment and processed with the sorting and multiplexed sequency strategy described previously (Figure 2A). A representative region of human chromosome 2, 57 to 122 Mb in human genome assembly 38, was selected to show the pattern of 50-kb-binned RTI calculated and smoothed via non-parametric LOESS regression. RPE p53KO MCM5^WT^ in dark blue is used as reference for pairwise comparison of the RTI profiles for ease of reading. Left side depicts results of our 2C-RT strategy based on Cyclin A2 staining, whereas right side shows data of traditional Repli-seq strategy based on DAPI staining. (G) Violin/box plots of delta RTI in untreated versus XL413 15 μM 4-hour treatment per 50-kb bin of each cell line showed in (F). Discontinuous line marks location for null RT difference, data below has lower RT values and data above has higher RT values in untreated condition. In each box, the line within the box indicates the median, and the box edges show the 25th and 75th percentiles. Whiskers extend beyond box limits 1.5 times the interquartile range between 25th and 75th percentiles. Dots indicate outliers past the whisker range.

Functionally, MCM5^bob1^ RPE cells maintained the proportion of mitotic cells following CDC7 inhibition while parental controls showed a strong reduction in mitotic index (Figure 3B). MCM5^bob1^ RPE cells also displayed fewer mitotic abnormalities than parental cells (Figure 3C and Figure S3A), consistent with more efficient replication completion. These findings align with previous reports that CDC7 inhibition causes mitotic errors such as ultra-fine anaphase bridges (Martin et al., 2022) and Mcm5-bob1 increases growth rate in *S. cerevisiae* only in the absence of Cdc7 (Hardy et al., 1997).

The bob1 mutation is predicted to allow the MCM hexamer to adopt conformations that can support origin activation even when replication origin priming is compromised (Hoang et al., 2007). Using 3D-SPARK, a super-resolution assay that employs sequential nascent DNA pulse labeling to detect and classify replication nanostructures (Hawgood et al., 2025), we quantified *initiation events* in individual cells. CDC7 inhibition significantly reduced the proportion of *initiation events* in parental cells but not in MCM5^bob1^ cells, supporting a model where MCM5^bob1^ partially bypasses CDC7 for origin activation (Figure 3D-E). Interestingly, flow cytometry revealed that overall DNA synthesis remained sensitive to CDC7 inhibition in MCM5^bob1^ cells, whereas RIF1KO cells maintained high EdU incorporation (Figure S3). These results suggest that RIF1 loss and MCM5^bob1^ mutation counteract CDC7 inhibition through distinct mechanisms and/or genomic contexts.

We performed 2C-RT and classic RT profiling in RIF1KO and MCM5^bob1^ cells in parallel to the parental p53KO controls described before (Figure 3F and Figure S4A). Under unperturbed conditions, MCM5^bob1^ and parental cells showed nearly identical RT profiles, with high inter-sample correlation across both methods (Pearson r = 0.96 in 2C-RT, 0.97 in classic RT). In contrast, RIF1KO cells exhibited pronounced RT changes compared to p53KO control and correspondingly lower correlation (Pearson r = 0.54 in 2C-RT, 0.54 in classic RT), consistent with previous reports (Yamazaki et al., 2012; Klein et al., 2021). To ensure that the differences between RIF1KO and p53KO cell lines were not dominated by the presence or absence of p53, we compared these data with published RT data from p53-proficient RPE cells generated by an independent laboratory (Van Rechem et al., 2021). The external dataset showed a high correlation with our p53KO data (Pearson r = 0.94 in 2C-RT, 0.96 in classic RT), suggesting that the observed differences are driven by RIF1 loss rather than p53 status (Figure S4B-D).

To quantify responses to CDC7 inhibition, we calculated RTI differences between mock and XL413 conditions (ΔRTI). All three cell lines exhibited delayed replication, but RIF1KO cells presented a less pronounced delay, consistent with a conserved role for RIF1 as a CDC7 antagonist. Although the range of ΔRTI values between mock and CDC7-inhibited MCM5^bob1^ cells was similar to that of parental cells, the distribution was shifted towards values closer to zero. This suggested that resistance to CDC7 inhibition in MCM5^bob1^ cells was not widespread but instead reflected more localized effects than those observed following RIF1 loss (Figure 3F-G).

The same comparison using the classic RT approach produced remarkably different results. According to DNA content, CDC7 inhibition caused a global RT advance in all three cell lines. In addition to being opposite in direction to the shift observed with 2C-RT, the effect was smaller in size and unable to differentiate between genotypes (Figure 3G). Taken together, these findings indicate that 2C-RT provides improved sensitivity and directional accuracy, enabling detection of replication dynamics that are not resolved by classic RT approaches.

### 2C-RT reveals a general RT advance in RIF1KO cells exacerbated at the nuclear periphery

Previous reports have suggested a nearly random RT program in human RIF1 knockout clones, implying that RIF1 delays replication in early regions as much as advances it in late regions. To address this, we performed pairwise comparisons of 2C-RT and classic RTI values between RIF1KO and RIF1-proficient cells (Figure 4A). Both approaches revealed a relatively similar genomic distribution of RTI changes, with a triangular shape comprising three basic RT clusters: late S-phase regions in RIF1-proficient p53KO cells that remained relatively the latest in RIF1KO; a small cluster of mid/late S-phase regions in p53KO that advanced to early S-phase in RIF1KO, and a large central area of early/mid S-phase regions in p53KO that present an RT in between the other two clusters for RIF1KO. We first isolated the latest replicating and most advancing clusters in 2C-RT (gates #1 and #2 in Figure 4A, respectively) and assessed the positioning of the gated bins in the classic RT graph, confirming that the selected RT clusters in both approaches contained the same genomic regions (Figures S5B-C). We then investigated their chromatin characteristics using publicly available data of chromatin states in RPE cells (Van Rechem et al., 2021). The latest replicating cluster (gate #1) contained mostly constitutive heterochromatin regions in RPE WT cells, whereas the most advanced regions in RIF1KO (gate #2) were enriched for a relatively unexplored euchromatic state marked solely by H3K36^me2^ (Figure 4B). Despite these shared overall patterns, the absolute shifts in RTI values compared to control differed substantially between 2C-RT and the classic RT approach. Direct comparison of RIF1KO data in 2C-RT and classic RT showed a genome-wide shift of 0.16 RTI units towards earlier RTI values in 2C-RT (Figure S5A). This change in the RTI distribution has important consequences for the inferred direction of RT changes and their mechanistic interpretation. Classic RT suggested that heterochromatic late S-phase regions did not change RT in RIF1KO cells, implying that these domains escape RIF1 control. In contrast, 2C-RT indicated that these regions do advance in RT, but not as dramatically as other regions, suggesting that RIF1 has a more extensive role than previously anticipated but does not exert equal pressure to postpone replication throughout the genome.

**Figure 4.**
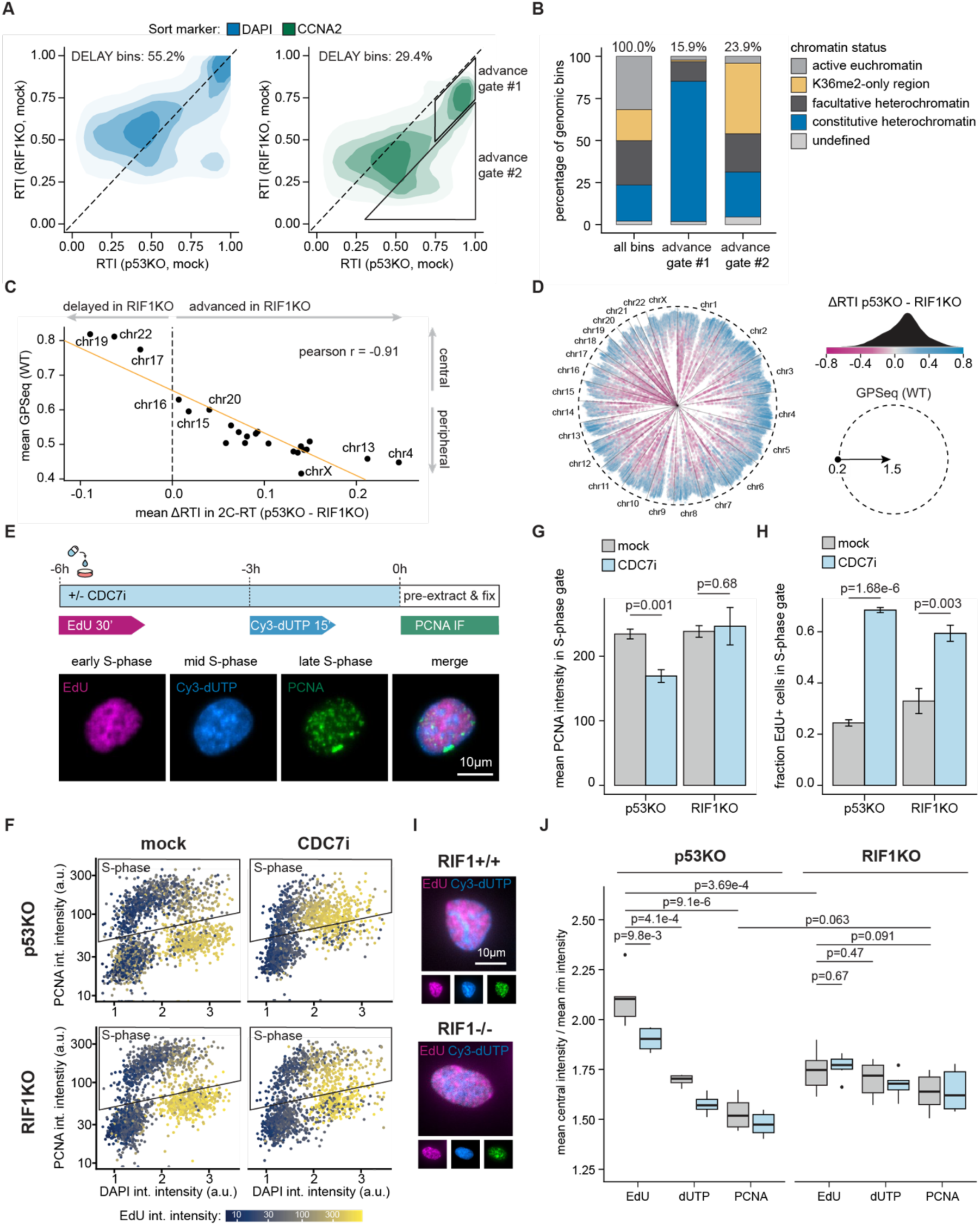
RIF1 loss globally accelerates RT with a more drastic shift in specific regions. A) Pairwise comparison of each 50-kb bin RTI value in p53KO RPE cells versus RIF1KO RPE cells. Each strategy is represented separately (2C-RT on the left, traditional Repli-seq on the right). Each contour area represents (from lighter to darker) 95%, 75%, 50% and 25% of the most densely regions covered by datapoints in the graph. Diagonal black line represents the expected location of a perfectly identical RT profile between condition. Percentage of 50-kb bins above the diagonal indicated at the top of each graph as “DELAY”. Gates #1 and #2 of genomic regions of interest are highlighted in black thick contour. B) Stacked bar plot of chromatin status in 50-kb genomic regions, either all regions or isolated in the specified gates of A). C) Dot plot of mean ΔRTI (p53KO – RIF1KO) of 2C-RT data per chromosome against mean GPSeq WT RPE cells per chromosome. Dashed vertical line at 0 marks the split between delay (lower than 0) and accelerated (higher than 0). Some extreme chromosome cases are indicated by name. Pearson correlation indicated at the top right. D) ΔRTI (p53KO – RIF1KO) in 50-kb bins radially arranged based on their GPSeq score and chromosome location (left), schematic of GPSeq score radial axis (right, bottom) and density plot of ΔRTI (p53KO – RIF1KO) matching each value with color within the diverging color palette (right, top) E) Schematic of the dual pulse-chase experiment performed on RPE p53KO or RIF1KO cells to assess DNA replication patterns at 6 hours (EdU), 3 hours (Cy3-x-dUTP) or right before fixation (PCNA). F) Dot plots of DAPI integrated intensity versus PCNA integrated intensity for RPE p53KO or RIF1KO underwent the dual pulse-chase experiment from E) with no drug or XL413 15 μM (CDC7i) for 6 hours. Dot colors match EdU integrated intensity. Representative images of each cell line, untreated on the right. G) Bar plot of mean value of PCNA integrated intensity in PCNA-positive cell population for RPE p53KO or RIF1KO cells that underwent dual pulse-chase experiment from E) with no drug or XL413 15 μM (CDC7i) for 6 hours (N = 6). Error bars indicate standard deviation. Statistical p-values of t-tests indicated between corresponding conditions compared. H) Bar plot of fraction of EdU-positive cells that are also PCNA-positive in RPE p53KO or RIF1KO cells that underwent dual pulse-chase experiment from E) with no drug or XL413 15 μM (CDC7i) for 6 hours (N = 6). Error bars indicate standard deviation. Statistical p-values of t-tests indicated between corresponding conditions compared. I) Box plot of the ratio of mean intensity at the inner part of the nucleus compared to mean intensity at the periphery of the same nucleus. Only triple-positive RPE p53KO or RIF1KO that underwent the dual pulse-chase experiment from E) with no drug or XL413 15 μM (CDC7i) for 6 hours were selected (N = 285 for RPE p53KO mock, N =852 for RPE p53KO CDC7i, N = 325 for RPE RIF1KO mock, N = 586 for RPE RIF1KO CDC7i). In each box, the line within the box indicates the median, and the box edges show the 25th and 75th percentiles. Whiskers extend beyond box limits 1.5 times the interquartile range between 25th and 75th percentiles. Statistical p-values of t-tests indicated between corresponding conditions compared.

Classic RT further suggested that nearly half of the genome is delayed in RIF1KO cells, whereas 2C-RT reported only a minor delay and mainly in early S-phase regions. This minor delay may reflect compensatory effects arising from the redistribution of limiting origin firing factors towards advancing regions (Zhu et al., 2025). We also evaluated the genome-wide RTI distribution in RIF1KO cells following CDC7 inhibition (Figure S5C). In accordance with our previous analysis (Figure 3G), classic RT showed no substantial change upon CDC7 inhibition regardless of RIF1 status (Figure 2H and S5C, blue graphs), while 2C-RT revealed a widespread replication delay smaller in RIF1KO cells than RIF1-proficient cells (Figure 2H and S5C, green graphs). This RTI pattern suggests that RIF1 preferentially targets specific genomic regions, whereas CDC7 acts with relatively uniform penetrance across the genome, consistent with a mobile nucleoplasmic kinase acting on an RT program already established on chromatin. It also reinforces that 2C-RT detects major shifts in DNA replication dynamics that remain unseen by classic RT methods.

Following previous reports of RIF1 localization at the nuclear envelope (Foti et al., 2016; Richards et al., 2022), we compared RTI shifts between p53KO and RIF1KO cells to the predicted nuclear position of the corresponding chromosomal territories based on published GPSeq data in RPE wild-type cells. Both 2C-RT and classic RT showed a strong correlation between RTI changes and proximity to nuclear envelope, in line with RIF1 association to the nuclear lamina (Figure 4C-D and S5D). We also examined the relationship between RT changes in RIF1KO cells and gene density, a chromosomal feature known to influence RT regulation. Chromosomes exhibiting the largest mean RT advances tended to have low gene density (Figure S5E).

When we evaluated the magnitude of RT changes, 2C-RT indicated most chromosomes experienced a noticeable RT advance in the absence of RIF1. In contrast, classic RT produced a mostly static picture, showing little change in mean RT for most chromosomes, and the ones that changed showed mostly delays (Figure S5D-E). This would imply that RIF1 plays a negligible role in regulating RT with respect to chromosome positioning. The chromosomes most affected by RIF1 loss tend to localize centrally and display relatively high gene density, which is inconsistent with published imaging and biochemical data describing predominantly peripheral RIF1 localization (Foti et al., 2016; Yamazaki et al., 2012; Zhu et al., 2025).

To complement our sequencing data and provide a different angle to the divergent results obtained with the two RT methods, we performed sequential nascent DNA pulse-chase experiments. Cells were incubated with EdU for 30 minutes at 6 hours before fixation and transfected with Cy3-x-dUTP 3 hours before fixation. Cells were either mock-treated or treated with XL413 at 6 hours before fixation and stained for chromatin-bound PCNA to define the S-phase population and measure replication proficiency at the end of the treatment (Figure 4E). RIF1KO cells displayed several features of increased replication efficiency under CDC7 inhibition, including higher levels of chromatin-bound PCNA in S-phase and a reduced fraction of EdU-positive cells within the S-phase population compared to parental cells (Figure 4F-H). We selected cells positive for EdU, dUTP and PCNA labelling and calculated center-to-rim intensity ratios for each channel (Figure 4I). In line with established RT patterns, RIF1-proficient cells showed minimal DNA synthesis activity incorporation at the nuclear periphery with the first pulse of EdU (reflecting early S-phase foci 6 hours before fixation), while the periphery showed stronger PCNA signal (reflecting mid/late S-phase foci at fixation). RIF1KO cells, however, displayed altered RT patterns: EdU was incorporated relatively more peripheral and PCNA localized more central in p53KO cells than in RIF1-proficient controls (Figure 4J). CDC7 inhibition did not overtly change spatial replication patterns in either RIF1KO or p53KO cells, consistent with the RT program not being primarily controlled at the kinase level. Together, these data support a model in which RIF1 does not randomize RT but instead delays replication onset in defined, predominantly peripheral chromosomal regions, thereby contributing to characteristic RT and replication foci patterns in S-phase.

### 2C-RT uncovers genetic and temporal control of initiation site preference

While examining RT profiles in MCM5^bob1^ cells, we noticed an unusual peak configuration. In XL413-treated MCM5^bob1^ cells, mid S-phase frequently showed clusters of EdU peaks of similar size, a pattern rarely observed under other conditions (Figure 5A, asterisks). These clusters of equally sized EdU peaks were most obvious at sites that displayed a single, well-defined early S-phase peak under untreated conditions, suggesting that this pattern reflects a response to CDC7 inhibition at early initiation sites.

**Figure 5.**
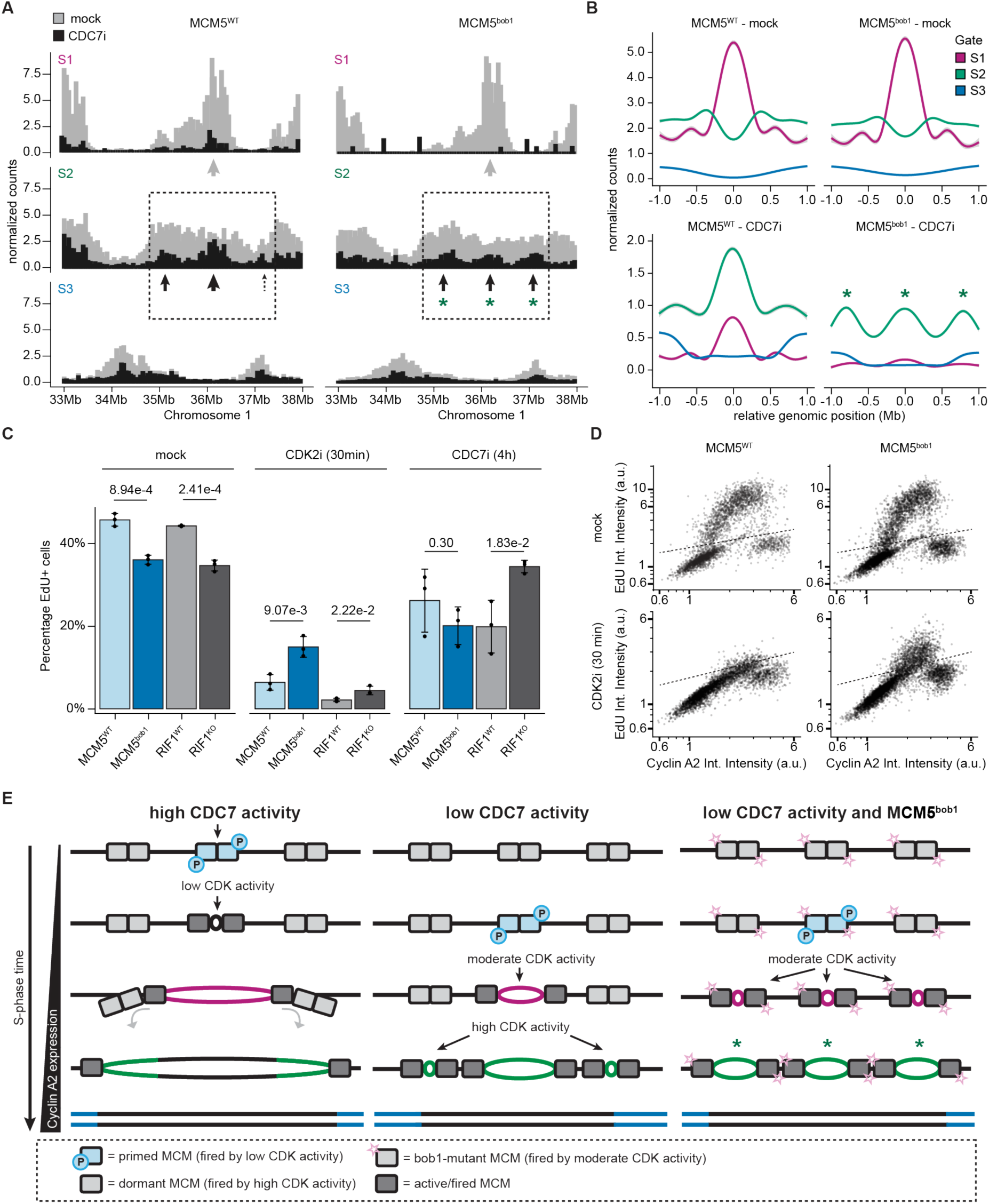
bob1 mutation presents a preferential switch towards CDK2-mediated firing over DDK-dependent origin activation. A) A representative region of human chromosome 1, 33 to 38 Mb in human genome assembly 38, showing an early initiation zone with the input-normalized 50-kb-binned 2C-RT tracks in each gate. Left side shows RPE p53KO MCM5^WT^ and right side MCM5^bob1^. Red box highlights the early initiation zone for mock, arrows mark the main peaks up to 1Mb away from initiation zone for XL413-treated cells. B) Aggregation line plots of 2C-RT normalized reads per gate for early initiation zones (N = 304). C) Bar plot of fraction of EdU-positive cells in RPE p53KO cells with wild-type MCM5 (MCM5^WT^), RPE p53KO cells with bob1-mutant MCM5 (MCM5^bob1^), RPE p53-proficient cells with functional RIF1 (RIF1^WT^) and RPE p53-proficient cells with no RIF1 expresion (RIF1^KO^) that were exposed for 30 minutes to NU6140 4μM (CDK2i) or for 4 hours to XL413 15 μM (CDC7i) (N = 3). Error bars indicate standard deviation. Statistical p-values of t-tests indicated between corresponding conditions compared. D) Dot plots of exemplary replicates quantified in 5C. Dotted line indicates cut-off of EdU vs cyclin A2 integrated intensity signals to identify actively replicating cells. E) Model for MCM5bob1 interaction with DDK-driven and CDK-driven origin priming programs for subsequent origin activation. Colors in the DNA tracks match the gate color scheme employed previously (red for S1, green for S2, blue for S3).

To systematically investigate this phenomenon, we analyzed 2C-RT read tracks for each gate to identify 304 early initiation sites in control cells (defined by EdU reads mainly in the S1 gate) and plotted normalized EdU signals within 1 Mb of the initiation centers (Figure 5B). In untreated RPE p53KO cells, regardless of MCM5 status, early initiation sites displayed a single dominant central peak in the S1 fraction that transitioned into a low-read valley flanked by broader peaks on each side in the S2 and S3 fractions, consistent with the expected progression of bidirectional DNA replication.

Upon CDC7 inhibition, however, the same initiation sites exhibited a different replication pattern depending on MCM5 status. In control cells, the central EdU peak was detected in both the S1 and S2 fractions, confirming a delay in replication initiation alongside persistent use of the dominant initiation site. In MCM5^bob1^ cells, by contrast, the central peak was flanked by two equally sized peaks approximately 750 kb away, suggesting a complete loss of initiation site dominance (Figure 5B). Interestingly, the response of MCM5^bob1^ cells to CDC7 inhibition changed as cells progressed through S-phase. EdU signal was nearly absent from the S1 fraction, indicating that replication was initially delayed. Nonetheless, genome-wide RTI profiles showed that replication was relatively advanced in XL413-treated MCM5^bob1^cells compared to parental controls, and mitotic fidelity was improved, consistent with DNA synthesis in mid or late S-phase compensating for the initial delay (Figure 3F-G). Supporting this interpretation, pairwise comparison of RTI in XL413-treated MCM5^wt^and MCM5^bob1^ cells showed that early S-phase regions were more delayed in MCM5^bob1^ cells, whereas mid/late S-phase regions were less delayed (Figure S6A-B)

Recent studies have provided evidence for two potential routes for origin firing: one relying on CDC7-dependent priming and low CDK2 activity, and one that is independent of CDC7 but requires high CDK2 activity (Suski et al., 2022; Upadhya et al., 2026). Given that Cyclin A2 levels rise during S-phase (Spencer et al., 2013; Akopyan et al., 2014) and directly promotes CDK2 activity during this phase (Merrick et al., 2008; Bendris et al., 2011), we hypothesized that the bob1 mutation promotes DNA replication in mid S-phase by facilitating CDK2-driven origin firing. To test this hypothesis, we treated MCM5^wt^ and MCM5^bob1^ cells with a short 30-minute pulse of CDK2 inhibitor NU6140 and quantified DNA synthesis and Cyclin A2 expression by QIBC (Fig 5C-D). The proportion of actively replicating (EdU^+^) cells was significantly higher in NU6140-treated MCM5^bob1^ cells compared to MCM5^wt^ controls and resistance was most evident in mid/late S-phase cells (as determined by high Cyclin A2 levels). These results support the idea that the bob1 mutation promotes DNA synthesis in mid/late S-phase by lowering the CDK2 activity threshold (Fig 5C-D). Interestingly, the bob1 mutation did not significantly affect the proportion of actively replicating cells upon CDC7 inhibition, indicating that the mutation does not alter the CDC7 activity threshold for replication initiation. In contrast, RIF1KO cells showed weaker resistance to NU6140 than MCM5^bob1^ cells but strong resistance to CDC7 inhibition, consistent with a role for RIF1 in controlling the CDC7 activity threshold via recruitment of the counteracting phosphatase.

Based on these observations, we propose a model for how the MCM5^bob1^ mutation attenuates the response to CDC7 inhibition while maintaining the global RT program (Figure 5E): CDC7 kinase primes origins within selected initiation zones, allowing these origins to be activated by low CDK1/CDK2 activity in early S-phase, regardless of the MCM5^bob1^ mutation. The bob1 mutation lowers the CDK1/CDK2 activity threshold for replication initiation, but before this threshold is reached, the majority of primed origins have fired; consequently, the effect of MCM5^bob1^ on RT is not apparent under unperturbed conditions. When CDC7 activity is low (e.g. by selective chemical inhibition), DNA replication is delayed genome-wide but the relative order and spatial regulation of RT are preserved. The initiation zones remain prioritized because residual CDC7 activity is still sufficient to prime origins and promote preferential firing in the same early-replicating regions before the alternative, CDC7-independent firing program driven by high CDK1/CDK2 activity takes over. The MCM5^bob1^ mutation lowers the threshold for activation of this alternative firing program. Thus, when replication onset is delayed in MCM5^bob1^ cells, the CDC7-dependent priming step is bypassed and all mutant MCM complexes exhibit a similar propensity for activation by CDK1/CDK2. The similar firing activity observed in initiation zones and proximal regions in CDC7-inhibited MCM5^bob1^ cells implies that the CDC7-independent program is less selective than the CDC7-primed firing program. RIF1 regulates RT by counteracting CDC7-mediated priming in defined chromatin environments, particularly in gene-sparse, peripheral chromosomal regions.

## Discussion

2C-RT provides a conceptual and technical advance that addresses a long-standing limitation in RT analysis. By uncoupling temporal inference from DNA content, it enables direct measurement of RT under conditions in which DNA synthesis is perturbed. Using Cyclin A2 as a DNA content–independent timer of S-phase progression, 2C-RT reveals genome-wide delays in replication following CDC7 inhibition that are effectively invisible to classic DAPI-based methods. These findings highlight how prior reliance on DNA content has likely obscured fundamental aspects of DNA replication dynamics.

This revised framework clarifies several paradoxes in the field, most notably the roles of RIF1 and CDC7. Rather than globally neutralizing RT, as previously proposed (Yamazaki et al., 2012; Klein et al., 2021; Caballero et al., 2023), RIF1 loss predominantly advances replication at specific loci, consistent with a model in which RIF1 spatially restricts CDC7 activity, particularly in peripheral or late-replicating chromatin environments (Hiraga et al., 2014; Alver et al., 2017; Vouzas & Gilbert, 2021). It further supports a model in which RIF1 exerts widespread but spatially localized control over RT (Figure 4A-D). In contrast, direct CDC7 inhibition with a small-molecule inhibitor causes a uniform RT delay across the genome (Figure 2H), suggesting that differential origin activity is regulated primarily at the RIF1–phosphatase level rather than at the kinase level.

2C-RT also separates global RT shifts from local origin activity regulation, as illustrated in the analysis of the MCM5^bob1^ bypass. The bob1 mutation is evolutionarily conserved and has minimal effect on RT in both yeast and human cells when CDC7 activity stays at regular levels (Hardy et al., 1997). However, when CDC7 activity becomes limiting, the intrinsically higher firing propensity of MCM5^bob1^ complexes through alternative mechanisms becomes apparent. This reduces origin dominance and mitigates mitotic chromosome segregation defects (Figure 3C and 5B). In yeast, this bypass remains dependent on S-phase cyclin activity, consistent with a model in which CDK input can sustain origin firing when DDK function is compromised (Sclafani et al., 2002; Hoang et al., 2007). In human cells, the preservation of the global RT program despite CDC7 inhibition or MCM5^bob1^ mutation suggests a similar principle, where CDK-supported initiation can maintain the overall temporal program, while CDC7 is specifically required to enforce origin selectivity within replication domains. The resulting phenotype is reminiscent of alternative initiation modes described in *Xenopus* extracts, in which late S-phase is associated with more frequent origin usage at the expense of slower fork progression (Ciardo et al., 2025). This similarity suggests that efficient bypass by MCM5^bob1^ may require a broader MCM activation regime.

Although replication initiation is built on a highly conserved ORC–MCM–CDC7 core, human cells operate within a more complex regulatory landscape that includes additional layers controlling licensing, firing, and stress responses, such as geminin, DONSON, RECQL4, and ETAA1 (Palm & Costa, 2025; Stillman et al., 2025; Yates et al., 2025). The ability to maintain a relatively intact RT program despite global perturbations, such as CDC7 inhibition or MCM5 mutation, implies that RT is maintained through multiple thresholds and feedback mechanisms (Upadhya et al., 2026). Our data support a layered model of RT control in which CDC7 primes early origins, CDK activity ensures genome-wide firing competence, and RIF1 imposes spatial constraints by counteracting CDC7 in specific genomic domains.

Continued development of perturbation-resilient RT profiling approaches such as 2C-RT, combined with targeted genetic models and high-resolution genomic and imaging techniques, will be essential for dissecting these interconnected mechanisms and linking RT changes to their spatial effectors. Even though established tools like Repli-Seq can offer similar data to 2C-RT from a relative sense (order of genomic regions replicated), it cannot properly establish its absolute timing position from S-phase onset. This is due to the direct relationship between DNA replication and DNA content change. DNA content as a metric of S-phase progression is confined to a range between the initial amount of DNA and double that amount, and can only change based on the ability of the cell to synthesize DNA at a given moment. For example, cells with half the genome duplicated will always show that half the genome is “early replicating” and the other half “late replicating” regardless of how long it took them to synthesize that much DNA. Consequently, DNA content-based approaches to infer replication timing thus give the impression of a globally stable RT program centered around mid S-phase (mean RTI = 0.5).

Counterintuitively, DNA content-based RT profiling methods compensate for drastic RT fluctuations in certain regions by shifting the rest of the regions in the opposite direction. This explains why CDC7 inhibition appears to produce a modest RT advance rather than the expected global delay. In the absence of CDC7, cells cannot start replicating DNA at the usual time from S-phase onset and have to wait for high CDK1/2 activity during mid/late S-phase in order to fire origins (Upadhya et al., 2026). This fail-safe mechanism may be less selective, as our MCM5^bob1^ data suggests and previous work describing a model in which late-replicating regions use numerous, slower forks (Ciardo et al., 2025; de los Rios Barreda et al., 2026). If cells halt early CDC7-dependent firing, it dramatically slows down DNA accumulation rate, “freezing” the timer until the subsequent firing program kicks in. Replication firing then proceeds at a faster pace in an unusually underreplicated genome, thus resuming DNA content build-up without an indication of a delay gap and giving the impression that normally late-replicating regions have advanced their timing. RT data from popular techniques like Repli-seq or EdU-seq therefore remain highly informative but should be carefully evaluated together with how DNA accumulation rate changes between the compared conditions.

Relative time estimates are now widely used in molecular biology, especially in single-cell omics, to order cells and infer dynamic processes from static measurements (Graham & Sottoriva, 2017; Ding et al., 2021). However, many of these approaches rely on correlations learned under steady-state or control conditions, which may not hold following perturbation. The risk of extrapolating such assumptions may apply when replication dynamics, transcriptional programs, RNA processing, or evolutionary rates are altered by stress, mutation, or regulatory changes (Caballero et al., 2023; Dietzen et al., 2024; de los Rios Barreda et al., 2026). Our findings highlight a broader opportunity in biological time inference, where perturbation-resilient clocks can substantially improve the interpretability of drug responses and dynamic developmental processes.

## Methods

### Cell culture

All cell lines were cultured in an ambient-controlled incubator at 37°C and 5% CO2. Human hTERT-immortalized retinal pigment epithelial cells (hereafter referred to as RPE) were obtained from the American Type Culture Collection (ATCC, CRL-4000) and maintained using DMEM-F12 GlutaMAX (Thermo Fisher Scientific, 31331093) supplemented with 1% penicillin-streptomycin (Thermo Fisher Scientific, 15140122) and 10% heat-inactivated fetal bovine serum (Sigma-Aldrich, F7524). The p53-deficient and isogenic parental RPE1 cells, previously described (Cascales et al., 2021), were kindly provided by Dr. Arne Lindqvist (KI). Homozygous bob1-mutant MCM5 RPE cells were generated for this study. The RIF1-deficient RPE cells previously described (Chiang et al., 2016) were generously provided by Dr. Steven P. Jackson (University of Cambridge). Human osteosarcoma U2OS cells were provided by Dr. Jiri Bartek (KI). Early passage skin fibroblast BJ cells were purchased from ATCC (CRL-2522). U2OS and BJ cells were cultured using DMEM-GlutaMAX (Thermo Fisher Scientific, 10566016) supplemented with 1% penicillin-streptomycin (Thermo Fisher Scientific, 15140122) and 10% heat-inactivated fetal bovine serum (Sigma-Aldrich, F7524). All cell lines were authenticated using IF and biochemistry methods within the last 3 years and regularly confirmed to be mycoplasma-free.

### MCM5^bob1^ cell line generation

RPE1 *p53*^−/−^ MCM5 bob-1 cells were generated by CRISPR Cas9-mediated knock-in via marker-free co-selection (Agudelo et al., 2017). Two guide RNA sequences (gRNA #A, 5′-GCTTGTACAAGTAGTCGGCC-3′; gRNA #B, 5′-GGGGGAGTACTGGATTGAGG-3′) targeting sites proximal to the MCM5 bob1 mutation were cloned into PX458_Cas9_GFP (Addgene #48138) using BbsI. For the HDR template, a 430 bp double-stranded DNA fragment (gBlock, IDT) containing a centrally positioned bob1 mutation was cloned into the backbone of ATP1A1_plasmid_donor_RD (Addgene #86551) using unique KpnI and EcoRI sites. The C>T bob1 mutation creates a novel NheI restriction site and disrupts the PAM sequence upstream of gRNA #A. A silent G>C substitution was introduced to disrupt the PAM sequence upstream of gRNA #B and prevent donor cleavage by the Cas9–sgRNA complex. The resulting PX458_bob1_A, PX458_bob1_B, and MCM5_bob1_HR_donor plasmids were sequence-verified and co-transfected with ATP1A1_plasmid_donor_RD and eSpCas9(1.1)_No_FLAG_ATP1A1_G3 (Addgene #86611) using FuGENE 6.Four days post transection, gene-targeted RPE1 *p53*^−/−^ cells were selected using 2.5 µM Ouabain (Sigma Aldrich #4995) and clonal lines were verified using PCR and NheI digest. Genomic DNA was isolated using QuickExtract DNA Extraction Solution 1.0 (Nordic Biosite) and PCR amplified using nested forward primers, bob-1 F(ex) TCCTCTTCTTTGCACCCTTCC and bob-1 F(in) CTCCAGGGATGAACTCAAGCG, and nested reverse primers, bob-1 R(ex) CATCAGCTACCTCCTTGGCAG and bob-1 R(in) seq CTATTAGTAACTGGGGCAGAGC. Two homozygous RPE1 *p53*^−/−^ MCM5^bob1^ clones were identified by complete NheI digestion and validated by sanger sequencing. The well-characterized Q118R/N129D mutations in *ATP1A1* introduce minimal modifications to the human genome with no detectable effect on cell fitness or on ATP1A1 steady-state protein levels or pump activity (Agudelo et al., 2017; Levesque et al., 2022). Sequenced regions were aligned to the reference MCM5 exon 3 locus using the alignment tools in Benchling (Benchling [Biology Software], 2025)

### Multiple sequence alignment of MCM5 across species

MCM5 protein sequences were retrieved from UniProt and aligned in Jalview using the MUSCLE multiple-sequence alignment algorithm with the protein preset settings (Edgar, 2004; Waterhouse et al., 2009). The following UniProt entries were analysed: Q6NV07 (*Danio rerio*), Q21902 (*Caenorhabditis elegans*), Q54CP4 (*Dictyostelium discoideum*), P41389 (*Schizosaccharomyces pombe*), Q0V8B7 (*Bos taurus*), P49718 (*Mus musculus*), Q561P5 (*Xenopus tropicalis*), B8AEH3 (*Oryza sativa indica*), Q6KAJ4 (*Oryza sativa japonica*), P29496 (*Saccharomyces cerevisiae*), P33992 (*Homo sapiens*), O80786 (*Arabidopsis thaliana*), Q9VGW7 (*Drosophila melanogaster*), P55862 (*Xenopus laevis – isoform A*), Q6PCI7 (*Xenopus laevis – isoform B*).

### Quantitative image-based cytometry (QIBC)

RPE cells were seeded in 96-well imaging plates (Sigma-Aldrich, CLS3904-100EA) 24 h before treatment with CDC7 inhibitor XL413 (15 µM; Sigma-Aldrich, SML1401), CDK2 inhibitor NU6140 (4 µM; Sigma-Aldrich, SML1323) or an equivalent volume of cell culture medium (mock). For metabolic labelling of nascent DNA, 5-ethynyl-2′-deoxyuridine (EdU; Jena Bioscience, CLK-N001-100) was added to the culture medium at a final concentration of 10 µM for 30 min. For nucleotide-transfection-based labelling of nascent DNA, culture medium was replaced with tricine buffer prepared as previously described (Kužmová et al., 2021) and containing aminoallyl-dUTP-x-Cy3 (6 µM; APExBIO, B8821) and synthetic nucleoside triphosphate transporter 1 (SNTT1; 6 µM; Sigma-Aldrich, SCT064). The labelling mixture was incubated at room temperature for 15 min before use. Cells were incubated with the transfection buffer for 4 min, washed with cell-culture-grade DPBS (Sigma-Aldrich, D8537), and returned to regular culture medium for 11 min, corresponding to a total labelling period of 15 min. Cells were then washed with DPBS and returned to regular culture medium as indicated. Following treatment and/or labelling, cells were washed with PBS (Gibco, 18912014), fixed in 4% paraformaldehyde (Histolab, 02176) for 7 min, permeabilized with cold methanol (Sigma-Aldrich) for 2 min, washed with PBS, and blocked in 2% BSA (Sigma-Aldrich, A9418) in TBS-T [Tris-buffered saline supplemented with 0.1% Tween-20]. Where indicated, cells were pre-extracted for 90 s in ice-cold pre-extraction buffer [0.5% Triton X-100 in CSK buffer] before fixation. For EdU detection, fixed cells were washed with DPBS and subjected to click chemistry in 100 mM Tris base (Sigma-Aldrich, T6066-1KG), 1 mM CuSO₄ (Sigma-Aldrich, C1297), 100 mM ascorbic acid (Sigma-Aldrich, A4544), and Alexa Fluor 647 azide (Life Technologies, A10277) for 1 h at room temperature. Cells were incubated overnight at 4 °C with primary antibodies diluted in blocking buffer: anti-cyclin A2 (1:500; Abcam, ab137769), anti-geminin (1:500; Proteintech, 10802-1-AP), anti-phospho-TCTP (Ser46) (1:500; Cell Signaling Technology, 5251), or anti-PCNA (clone PC10; 1:400; Santa Cruz Biotechnology, sc-56). Samples were washed with TBS-T and DPBS, then incubated for 1 h at room temperature with DAPI and the appropriate secondary antibody diluted in blocking buffer: Alexa Fluor 488-conjugated anti-rabbit IgG (1:1,000; Life Technologies, A11008), Alexa Fluor 555-conjugated anti-rabbit IgG (1:1,000; Life Technologies, A21428), Alexa Fluor 488-conjugated anti-mouse IgG (1:1,000; Life Technologies, A11001), or Alexa Fluor 555-conjugated anti-mouse IgG (1:1,000; Life Technologies, A21422). Samples were stored in DPBS at 4 °C until imaging.

Widefield images were acquired at room temperature using a Nikon Eclipse Ti2 microscope equipped with either a 20× air objective or a 40× water-immersion objective. More than 1,000 cells were analyzed per replicate and condition. Nuclei were segmented using custom CellProfiler pipelines, and downstream analyses were performed in R. In asynchronous cell populations, cell-cycle phases (G1, S, and G2) were assigned using an active replication marker—EdU incorporation or nuclear PCNA intensity—in combination with an S-phase progression marker, DAPI signal or cyclin A2 abundance. For release time-course experiments, nuclear DAPI, EdU, cyclin A2, and geminin intensities were normalized to the residual G1/G0 population. The median normalized signal intensity of the released cell population was used to monitor cell-cycle progression over time. For analysis of mitotic errors, high phospho-TCTP signal was used to identify mitotic cells. Mitotic stage and the presence of anaphase bridges were subsequently verified by manual inspection of DAPI-stained nuclear structures. For pulse–chase–pulse analysis, S-phase cells were defined based on nuclear PCNA intensity. EdU-positive and Cy3-dUTP-positive cells were identified using thresholds established from the corresponding G1-cell populations. Nuclear-rim and nuclear-interior regions were generated by shrinking the nuclear masks by 3 pixels from the nuclear boundary.

### 3D Spatial Assay for Replication Kinetics (3D-SPARK)

3D-SPARK was performed as previously described (Hawgood et al., 2025). RPE cells were seeded on 18-mm high-precision no. 1.5 glass coverslips (Marienfeld, 0111580) and subjected to the standard pulsing scheme: a 15-min EdU pulse followed immediately by a 15-min aminoallyl-dUTP-x-Cy3 pulse (APExBIO, B8821). Cells were fixed in 4% paraformaldehyde, and EdU click chemistry and DAPI staining were performed as described above. Coverslips were mounted in ProLong Gold Antifade Mountant (Thermo Fisher Scientific, P10144) on SuperFrost Ultra Plus Adhesion Slides (Thermo Fisher Scientific, 1014356190), cured for 24 h in the dark at room temperature, and stored at 4 °C until imaging. Three-dimensional structured illumination microscopy (3D-SIM) was performed using a Carl Zeiss Elyra PS.1 microscope equipped with 405-, 488-, 561-, and 642-nm lasers. Images were acquired sequentially using an Andor iXon DU 897 camera and a Plan-Apochromat 100×/1.46 NA oil-immersion objective. Emission signals were collected through appropriate dichroic mirrors and band-pass filters (570–650 nm for 561-nm excitation and >655 nm for 642-nm excitation). For each image, the illumination grid was rotated through five angular positions and acquired at five phase shifts per angle. Raw 3D-SIM data were reconstructed using the integrated ELYRA S software (ZEN 2011 SP2 Black) with automated reconstruction settings. Calibration using 40-nm sub-resolution beads yielded a lateral resolution of approximately 85 ± 5 nm at 488-nm excitation, as described previously (Agostinho et al., 2016). Downstream segmentation of 3D-SIM-reconstructed nuclei was done in Python to quantify fraction of initiation events per cell. Analysis code available in GitHub repository (https://github.com/LemmensLab/3D-SPARK).

### Multiplexed nascent DNA sequencing

For multiplexed nascent DNA sequencing, we adapted the MINUTE-ChIP protocol (Kumar et al., 2025; Kumar & Elsässer, 2019) with several modifications to accommodate nascent DNA pulldown. RPE cells were seeded in 150-mm dishes at a density sufficient to recover 1 × 10⁷ cells per condition and sort strategy. Cells were pulse-labeled with 10 μM EdU prior to trypsinization, collected in culture medium in non-sterile 50 mL tubes, and fixed with 1% formaldehyde (Pierce™ 16% Formaldehyde, Thermo Fisher Scientific, #28908) for 10 min at room temperature with rotation. Residual reactive formaldehyde was quenched by adding Tris-HCl (3 M, pH 7.8; to a final concentration of 600 mM) for 5 min at room temperature with rotation. Fixed cells were permeabilized with 0.5% saponin (Sigma-Aldrich, #47036), 0.05% IgG-free BSA (Jackson ImmunoResearch, #001-000-161), and 1× Halt™ Protease and Phosphatase Inhibitor Cocktail (Thermo Fisher Scientific, #78444) for 30 min at 4°C with rotation, followed by a second 30-min incubation at 4°C with rotation in the same solution adjusted to 0.5% IgG-free BSA. Cell concentration was adjusted to 1–2 × 10⁷ cells/mL, and the EdU moiety was clicked for 30 min at room temperature in the dark using the following reaction mix: 50% DMSO, 0.05% saponin, 2 mM CuSO₄ (Sigma-Aldrich, C1297), 2 mM BTTAA (Jena Bioscience, CLK-067-100), 10 mM sodium ascorbate (Sigma-Aldrich, A7631), and 500 nM picolyl azide label (Alexa Fluor 488, Jena Bioscience, CLK-1276-AZ; or PEG4-biotin, Jena Bioscience, CLK-1167). The click reaction was quenched with EDTA (VWR, #1.08418.0100) at a final concentration of 20 mM for 5 min at room temperature. Cells were then incubated overnight at 4°C in the dark with primary antibody (anti-cyclin A2, 1:500, Proteintech, 66391-1-Ig; anti-mCherry, 1:200, Abcam, ab124765) in incubation solution (0.5% saponin, 0.05% IgG-free BSA, 2 mM EDTA, 1× protease and phosphatase inhibitor cocktail). Cells were washed twice in incubation solution without antibody and incubated for 2 h at 4°C in the dark with secondary antibody (rabbit IgG–Alexa Fluor 647, 1:500, Life Technologies, #A21245; or mouse IgG–Alexa Fluor 647, 1:500, Life Technologies, #A21235) and 1 μg/mL DAPI in incubation solution. Finally, cells were washed twice in incubation solution, with 15-min incubations at 4°C in the dark between washes to destain DAPI.

To collect the different cell cycle fractions, cell concentration was adjusted to 2 × 10⁷ cells/mL for each sample, and cells were sorted on a FACSAria Fusion (BD Biosciences) using four gates to capture EdU-negative G1 cells and EdU-positive cells with low (S1), intermediate (S2), or high (S3) DNA content or cyclin A2 levels. Individual paraformaldehyde-fixed, FACS-sorted, cell-cycle-phase-specific samples (100,000–250,000 cells) were MNase-digested to enrich for mononucleosomal fragments, which were ligated to sample-barcoding adapters from the EpiFinder™ GenomePro kit (Epigenica) following the manufacturer’s instructions. Uniquely barcoded samples were pooled, and crosslinks were reversed by overnight proteinase K digestion before purification with AMPure beads. Ten percent of the recovered DNA was reserved as input, and the remainder was subjected to DNA immunoprecipitation (DIP). Pooled samples containing EdU–Alexa Fluor 488 or EdU–biotin click conjugates were subjected to DIP using, respectively, Protein A beads coupled to anti-Alexa Fluor 488 antibody (Thermo Fisher Scientific, #710369) in PBS with 0.065% Triton X-100, or MyOne Streptavidin T1 Dynabeads in 10 mM Tris-HCl (pH 7.5), 1 mM EDTA (pH 8.0), 2 M NaCl, and 0.1% Tween-20. After washing in the respective DIP buffer, bead-enriched DNA was eluted by proteinase K treatment and purified with the Zymo Research DNA Clean & Concentrator kit. Libraries were constructed from the DIP-enriched and input DNA following the MINUTE-ChIP workflow, starting with in vitro transcription, RA3 3′ adapter ligation, cDNA conversion, and PCR amplification with barcoded primers compatible with the Illumina sequencing platform. Library fragment profiles were assessed on an Agilent Bioanalyzer, quantified with the Qubit dsDNA High Sensitivity assay, and diluted for sequencing on an Illumina NextSeq 2000.

FASTQ files from the sequencing runs were processed as described in the MINUTE-ChIP analysis protocol to obtain input-normalized reads mapped to the human genome assembly hg38 and demultiplexed per condition with a custom snakemake pipeline (https://github.com/elsasserlab/minute). Input-normalized reads were aggregated into 50-kb bins using the wigglescout package (https://github.com/cnluzon/wigglescout). A replication timing index (RTI) was calculated per bin and condition using the following formula (where n denotes the number of EdU-positive sorted gates per condition):

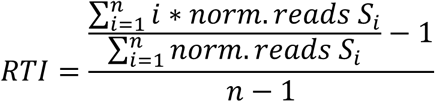

For Early initiation zone analysis, we filtered genomic regions in RPE p53KO (MCM5WT, mock) 2C-RT dataset with the following criteria: RTI of genomic region < 0.125 (>75% reads come from S1 gate) and normalized read count of genomic region in S1 is the local maxima in −/+ 1Mb range. Genomic coordinates of those regions were selected for further examination in the rest of the conditions.

### Comparison of 2C-RT results with public genomics data

Public datasets from p53-proficient hTERT RPE-1 cells were obtained for the following chromatin features: H3K9me3, RTI, chromatin state based on multiple epigenetic marks in assembly hg19 (Van Rechem et al., 2021); TSA-seq proximity to laminB1 in assembly hg19 (Chen et al., 2018); GPSeq nuclear positioning in assembly hg38 (Yip et al., 2026). Datasets mapped to genome assemblies other than hg38 were converted to hg38 coordinates using the UCSC LiftOver tool (Casper et al., 2026; Hinrichs et al., 2006) to roughly convert genome coordinates between assemblies. New coordinates were matched to overlapping 50-kb regions overlapping by more than 50% in our dataset for downstream analysis.

## Acknowledgements

We thank Ana Agostinho from the Advanced Light Microscopy Facility at KTH-SciLifeLab, and the National Microscopy Infrastructure NMI (VR-RFI 2023-00163 for 3D-SIM super-resolution imaging support. We are grateful to Dr. Magda Bienko, Wing Hin Yip and Kaja Harton for technical support and valuable feedback on GPSeq analysis, and Dr. Arne Lindqvist (Karolinska Institutet) for providing the RPE1 p53KO cell line and Dr. Steven P. Jackson (University of Cambridge) for gifting the RIF1-deficient RPE1 cells and parental controls. This research was made possible by an ASPIRE Award from The Mark Foundation for Cancer Research (Grant No. 23-040-ASP). We gratefully acknowledge funding from the Swedish Research Council (Starting grant 2019-04667; project grant VR-MH 2014-46602-117891-30), Karolinska Institutet (Assistant Professor Faculty grant 2-1534/2020; KID doctoral education grant; and SciLifeLab (RED project grant).

---

**Figure S1.**
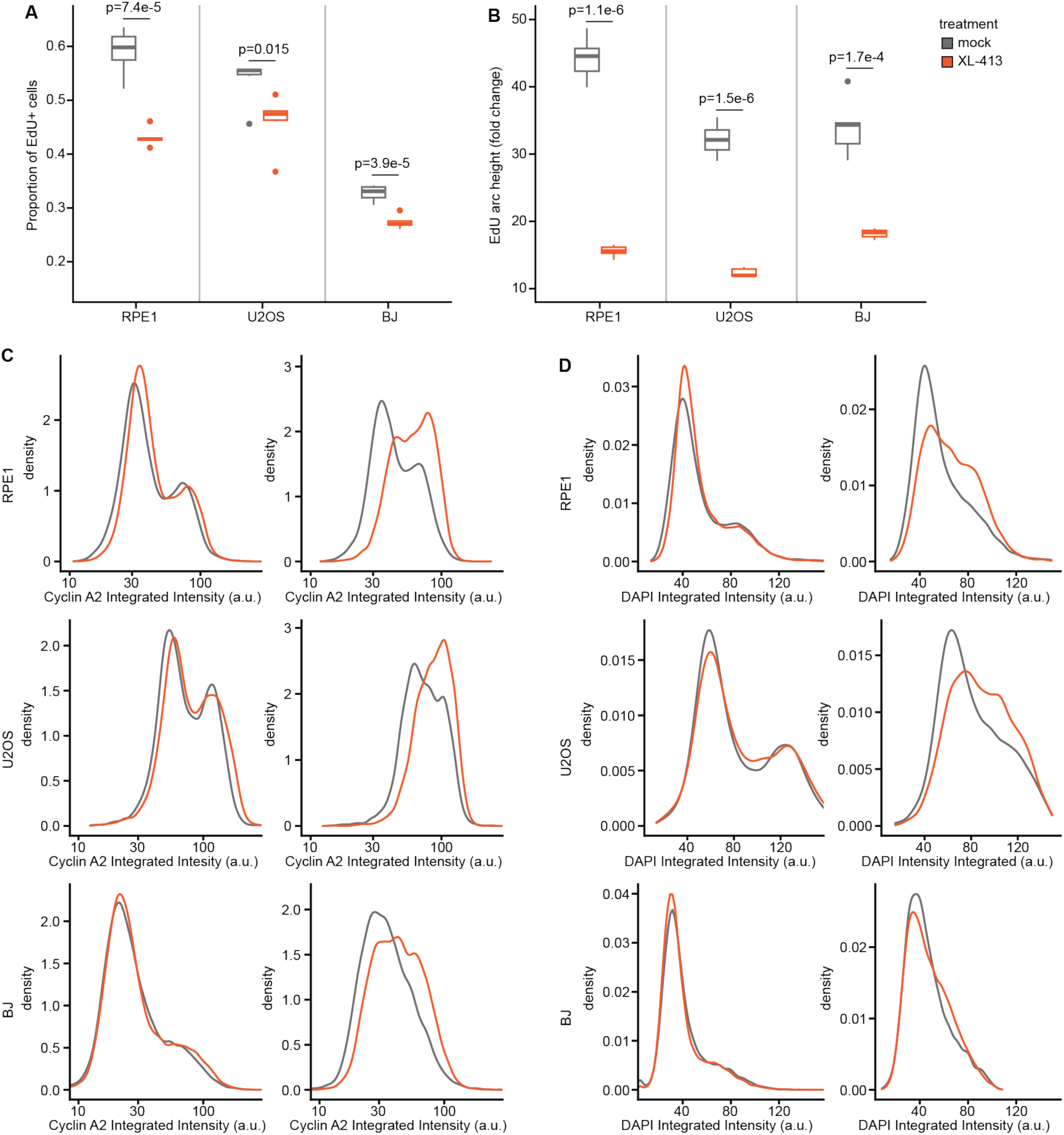
Differential trends of DNA content and Cyclin A2 accumulation upon CDC7 inhibition are constant in human cells. Different human cell lines (RPE1 and U2OS cells) were treated with no drug or XL413 15 μM for 4 hours and labelled for DNA synthesis in the last 30 minutes of treatment with EdU 10 μM prior to fixation. Cells were stained for DNA content or Cyclin A2. (A) Box plot of proportion of EdU-positive cells in each cell line for each treatment described before, N = 6. In each box, the line within the box indicates the median, and the box edges show the 25th and 75th percentiles. Whiskers extend beyond box limits 1.5 times the interquartile range between 25th and 75th percentiles. Statistical p-values of t-tests indicated between corresponding conditions compared. (B) Box plot of EdU fold-change from median integrated intensity signal in G1 population to 10% top percentile integrated intensity signal in S population for each cell line and drug treatment described before, N = 6. In each box, the line within the box indicates the median, and the box edges show the 25th and 75th percentiles. Whiskers extend beyond box limits 1.5 times the interquartile range between 25th and 75th percentiles. Statistical p-values of t-tests indicated between corresponding conditions compared. (C) Density plots of Cyclin A2 integrated intensity of whole population (left) or EdU-positive population (right) for each cell line and treatment as above. Including both whole and EdU-positive profiles allows better visual comparison between cell lines, otherwise differing immunostaining success would mask the true fluctuation of the EdU-positive population. (D) Same as (C) but plotting DAPI integrated intensity instead.

**Figure S2.**
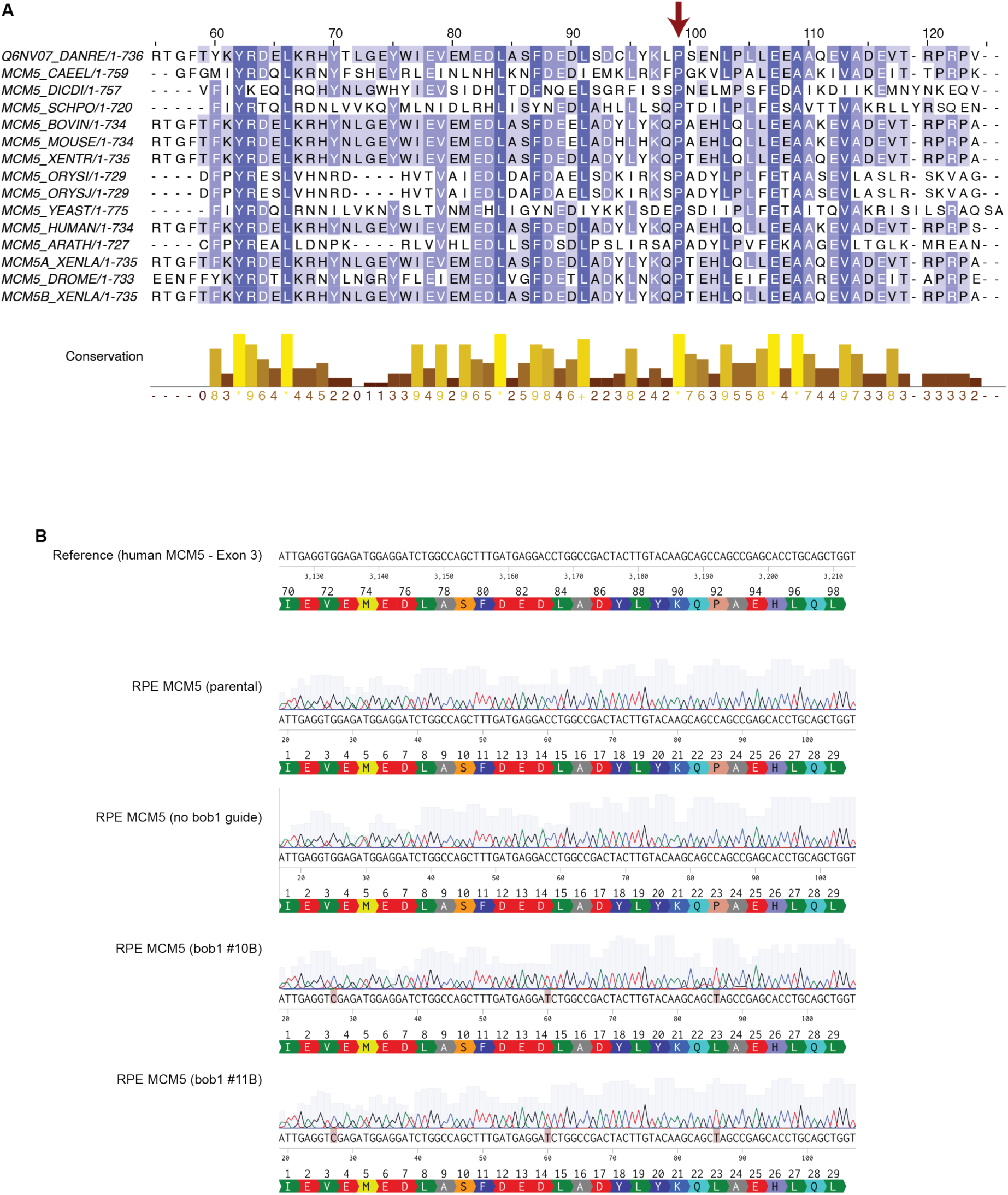
Validation of human bob1 knock-in. (A) Multiple sequence alignment of 15 different MCM5 protein sequences across different eukaryotes. Alignment was performed in Jalview 2.11.5.2 using MUSCLE preset for protein alignment. Results show positions 55 to 126 of the resulting alignment, which include proline 83 in *Saccharomyces cerevisiae*, highlighted with a dark red arrow. Residue conservation degree is indicated by darker blue per position on the alignment and also summarized at the bottom barplot. (B) Pairwise alignment of sequenced RPE cell lines used or generated during the MCM5^bob1^ CRISPR knock-in. The region spanning exon 3 of human MCM5 gene, where conserved proline 92 lies, was amplified and sequenced. Pairwise alignment against human MCM5 gene reference (ENST00000216122) was performed with Benchling electronic notebook, allowing simultaneous visualization of sequencing quality for homozygous knock-in, mutated sites to reference and resulting translational changes. Only RPE bob1 clones #10B and #11B showed homozygous knock-in of bob1 mutation and Cas9 PAM sites evaluated for knock-in.

**Figure S3.**
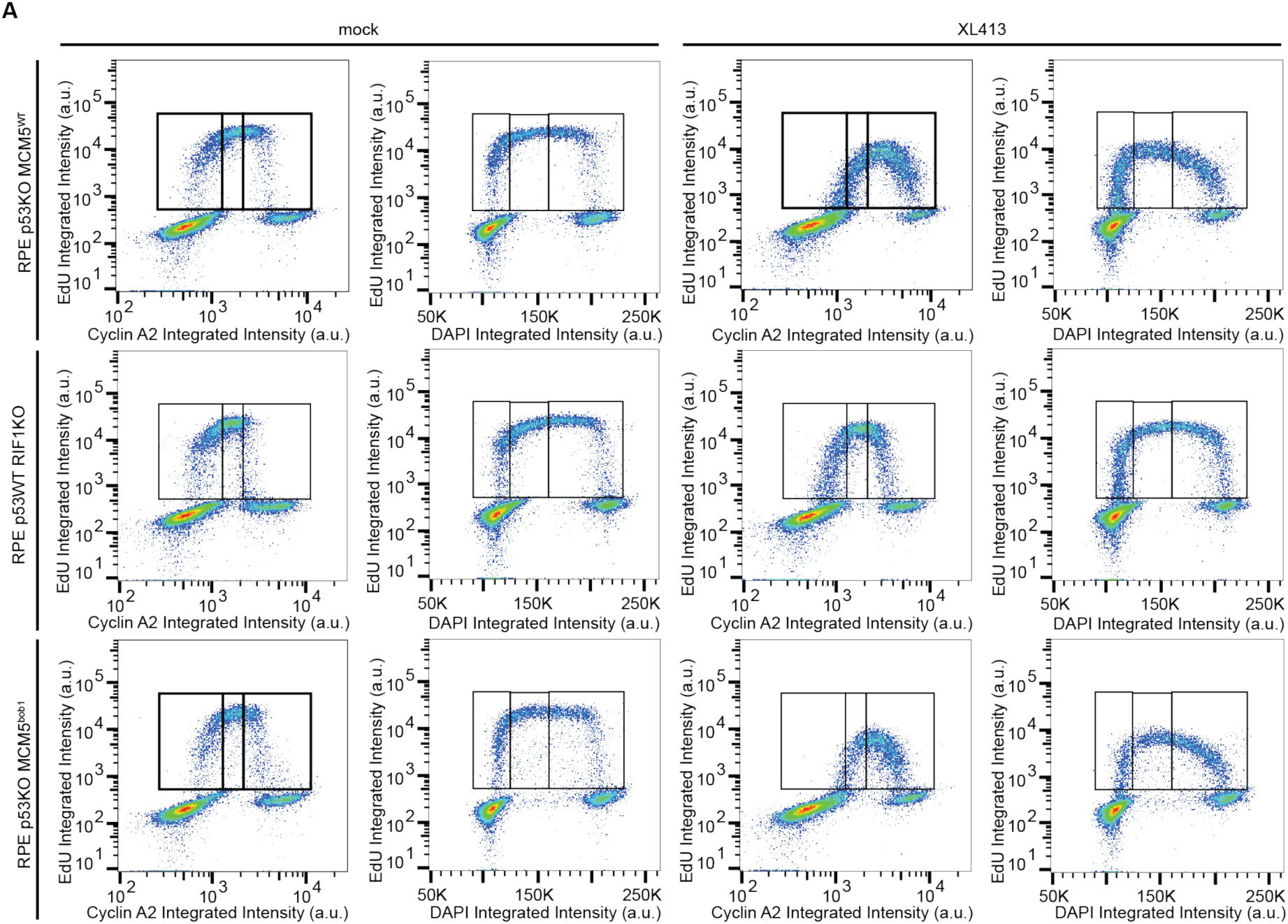
Cyclin A2 levels capture DNA replication delay upon CDC7 inhibition in sorted cells. (A) Flow cytometry data of the different RPE cell genotypes sorted for RT evaluation applying 2C-RT or traditional Repli-seq. All cells were treated with no drug or XL413 15 μM (Cdc7i) for 4 hours and labelled for nascent DNA with EdU 10 μM in the last 30 minutes prior to collection. The data for p53KO parental is the same as in Figure 1E, added here for easier comparison.

**Figure S4.**
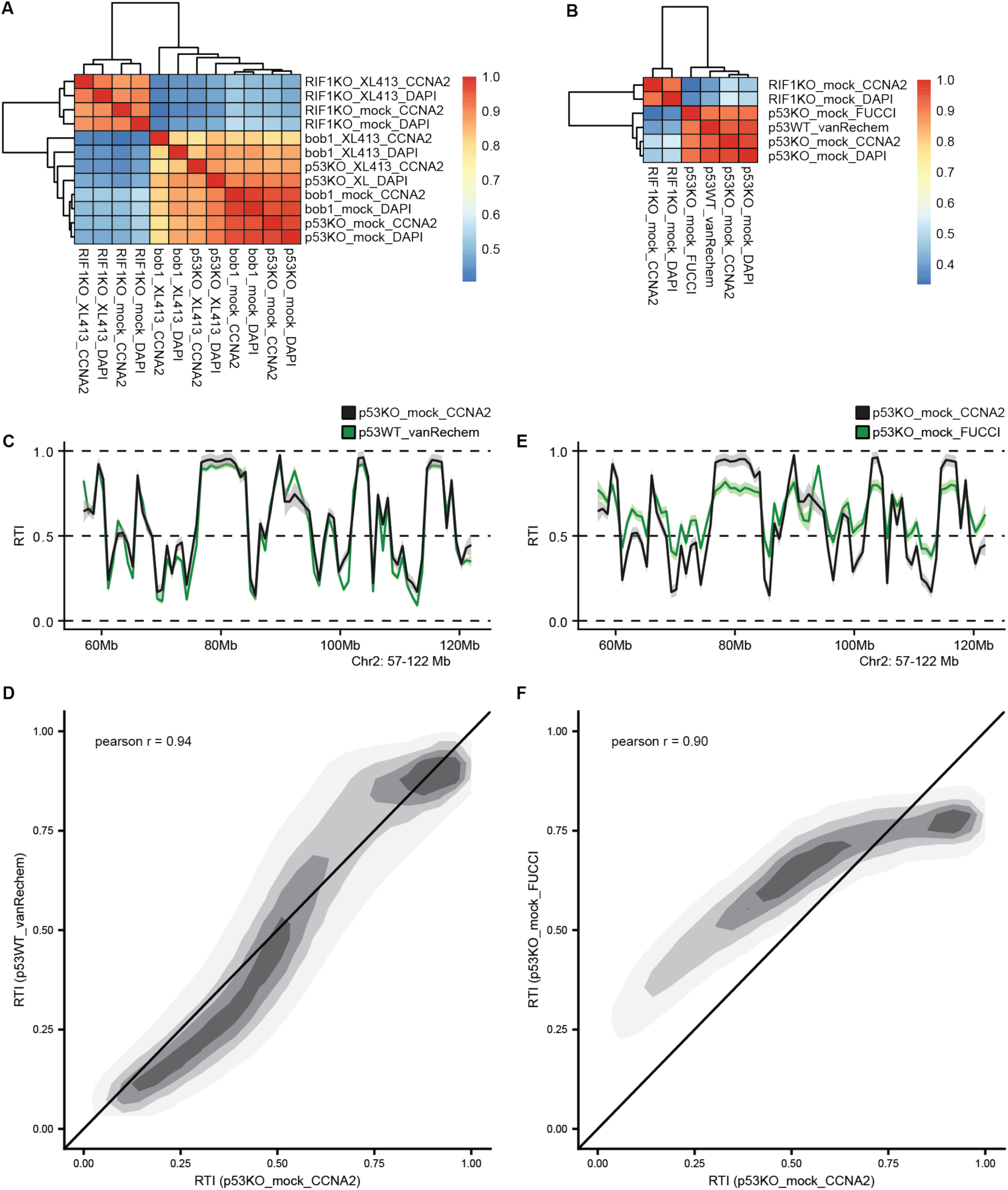
2C-RT and DNA-content-based RT strategies highly correlate in unperturbed condition. (A) Correlation matrix heatmap of the pearson correlations for every 50-kb binned RTI pairwise comparison of the different combinations of cell lines, treatments and sorting strategies implemented in Figure 3. Hierarchical clustering dendrograms of all conditions indicated at the top and left of the heatmap. 2C-RT data is indicated as “CCNA2”, as they are based on Cyclin A2 accumulation. (B) Correlation matrix heatmap of the pearson correlations for every 50-kb binned RTI pairwise comparison of the untreated RPE p53KO in 2C-RT and DNA-content-based, RPE RIF1KO in 2C-RT and DNA-content-based, published RTI data on RPE p53 wild-type (p53WT_vanRechem) and a sequencing run from RPE p53KO cells containing PIP-FUCCI system that we sorted based on ectopic Geminin_1-110_-mCherry levels (p53KO_mock_FUCCI). Hierarchical clustering dendrograms of all conditions indicated at the top and left of the heatmap. 2C-RT data is indicated as “CCNA2”, as they are based on Cyclin A2 accumulation. (C) RTI profile smoothed by LOESS non-parametric regression of 2C-RT untreated data in RPE p53KO and published Repli-seq data in RPE p53 wild-type. A representative region of human chromosome 2, 57 to 122 Mb in human genome assembly 38, was selected to show the pattern of 50-kb-binned RTI calculated. Lighter shadow represents 95% confidence interval of the regression. (D). (E) and (F) are equivalent to (C) and (D) respectively, but comparing 2C-RT results to FUCCI-based sorting on RPE p53KO cells instead.

**Figure S5.**
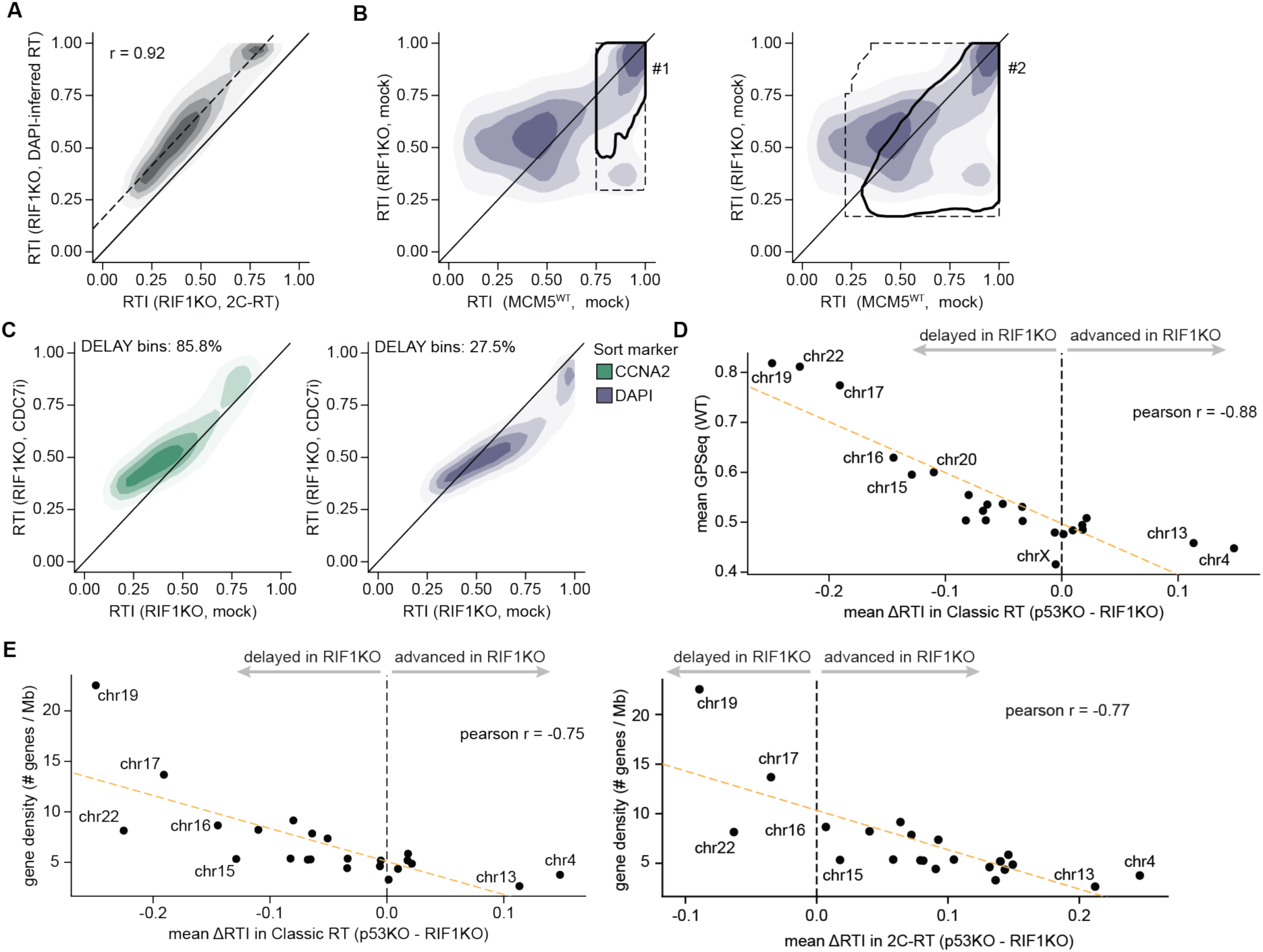
Classic RT keeps RT distribution balanced around mid-RT value range. A) Pairwise comparison of each 50-kb bin RTI value in RIF1KO untreated RPE cells obtained with 2C-RT against classic RT. Each contour area represents (from lighter to darker) 95%, 75%, 50% and 25% of the most densely regions covered by datapoints in the graph. Diagonal black line represents the expected location of a perfectly identical RT profile between condition. Dashed line indicates linear regression for the pairwise data. Pearson correlation value at the top left. B) Pairwise comparison of each 50-kb bin RTI value in p53KO RPE cells against RIF1KO RPE cells obtained with classic RT. Each contour area represents (from lighter to darker) 95%, 75%, 50% and 25% of the most densely regions covered by datapoints in the graph. Diagonal black line represents the expected location of a perfectly identical RT profile between conditions. Black dashed contour area indicates location of genomic bins within each gate highlighted in Figure 4A, 2C-RT dataset. Black solid contour area indicates location of 99% most clustered genomic bins in classic RT coming from each gate highlighted in Figure 4A, 2C-RT dataset. C) Pairwise comparison of each 50-kb bin RTI value in mock versus XL413-treated RPE RIF1KO cells. Each strategy is represented separately (2C-RT on the left, traditional Repli-seq on the right). Each contour area represents (from lighter to darker) 95%, 75%, 50% and 25% of the most densely regions covered by datapoints in the graph. Diagonal black line represents the expected location of a perfectly identical RT profile between condition. Percentage of 50-kb bins above the diagonal indicated at the top of each graph as “DELAY”. D) Dot plot of mean ΔRTI (p53KO – RIF1KO) of classic RT data per chromosome against mean GPSeq WT RPE cells per chromosome. Dashed vertical line at 0 marks the split between delay (lower than 0) and accelerated (higher than 0). Some extreme chromosome cases are indicated by name. Pearson correlation indicated at the top right. E) Dot plot of mean ΔRTI (p53KO – RIF1KO) of classic RT (left) or 2C-RT (right) data per chromosome against gene density per chromosome. Dashed vertical line at 0 marks the split between delay (lower than 0) and accelerated (higher than 0). Some extreme chromosome cases are indicated by name. Pearson correlation indicated at the top right.

**Figure S6.**
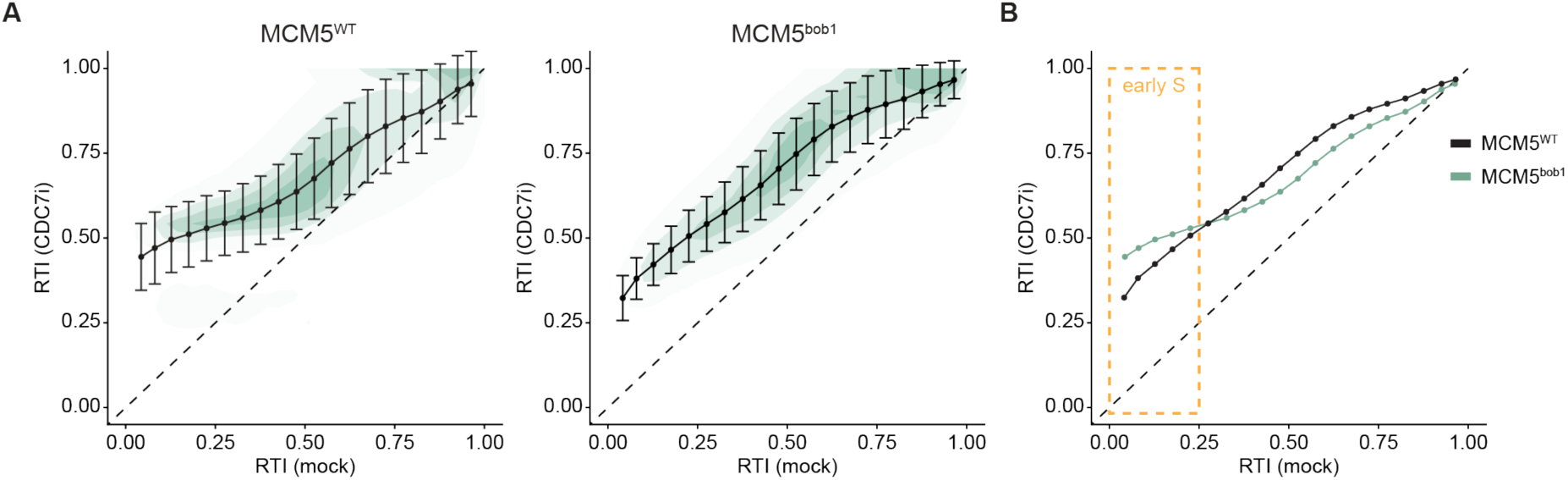
2C-RT shows compensatory effects of bob1 mutation against CDC7 inhibition in mid/late S-phase. A) Pairwise comparison of each 50-kb bin 2C-RT RTI value in mock versus XL413-treated of RPE p53KO MCM5 bob1-mutant cells (left) and RPE p53KO MCM5 WT cells (right, data as in Figure 2G). Each contour area represents (from lighter to darker) 95%, 75%, 50% and 25% of the most densely regions covered by datapoints in the graph. Bins were grouped by RTI value in mock condition, forming 20 contiguous groups of 0.05 RTI range each. Dots represent mean RTI value in XL413-treated condition for the group of genomic regions, while whiskers indicate one standard deviation of the corresponding group. Diagonal dashed line represents the expected location of a perfectly identical RT profile between conditions. B) Combination of graphs in (A), but only showing the mean RTI per 0.05 range group for direct comparison of the RT trends between cell lines. Yellow dashed box indicates the RTI values that we describe as “early” in mock conditions.

